# Probing the limits of activity-silent non-conscious working memory

**DOI:** 10.1101/379537

**Authors:** Darinka Trübutschek, Sébastien Marti, Henrik Ueberschär, Stanislas Dehaene

## Abstract

Two types of working memory (WM) have recently been proposed: conscious active WM, depending on sustained neural activity, and activity-silent WM, requiring neither conscious awareness nor accompanying neural activity. However, whether both states support identical forms of information processing is unknown. Theory predicts that activity-silent states are confined to passive storage and cannot operate on stored information. To determine whether an explicit reactivation is required prior to the manipulation of information in WM, we evaluated whether participants could mentally rotate brief visual stimuli of variable subjective visibility. Behaviorally, even for unseen targets, subjects reported the rotated location above chance after several seconds. As predicted, however, such blindsight performance was accompanied by neural signatures of conscious reactivation at the time of mental rotation, including a sustained desynchronization in alpha/beta frequency and a decodable representation of participants’ guess and response. Our findings challenge the concept of genuine non-conscious “working” memory, argue that activity-silent states merely support passive short-term memory, and provide a cautionary note for purely behavioral studies of non-conscious information processing.

## Introduction

Working memory (WM) serves a critical role in the online storage of information for rapid access, transformation, and flexible use. Until recently, it was thought to depend on conscious, effortful processing (Baars and Franklin, 2003; Baddeley, 2000, 2003) and the maintenance of persistent neural activity (Fuster and Alexander, 1971; Goldman-Rakic, 1995; Kamiński et al., 2017). However, a growing body of evidence suggests that successful WM maintenance may be dissociated from consciousness and persistent delay-period activity. Items subjectively reported as unseen may still be retrieved above chance-level after several seconds (Bergström and Eriksson, 2014, 2015; King et al., 2016; Soto et al., 2011; Trübutschek et al., 2017). Likewise, an uninterrupted chain of persistent neural firing is not always observed during WM maintenance (Watanabe and Funahashi, 2007, 2014) and content-specific delay-period activity may vanish during the maintenance of non-conscious or unattended information (Rose et al., 2016; Trübutschek et al., 2017; Wolff et al., 2015, 2017).

Theories and simulations indicate that such “activity-silent” maintenance in the absence of accompanying neural activity may be supported by short-term changes in synapses temporarily linking populations of neurons coding for the stored items (Mongillo et al., 2008; Stokes, 2015). Later, a non-specific stimulation of the system may reinstate the original neural firing pattern, an effect that was recently observed experimentally (Rose et al., 2016; Wolff et al., 2017). Short-term synaptic changes may thus effectively allow networks to go silent for several seconds while still supporting a delayed information readout.

While the evidence for active versus activity-silent forms of WM is mounting, whether they support identical forms of information processing remains unknown. Beyond maintenance, a defining feature of WM is the ability to manipulate information, for instance during mental rotation (Baddeley, 1992; Luck and Vogel, 2013). If non-conscious WM representations are indeed stored via activity-silent short-term synaptic changes, it is unclear whether they might be transformed without first being reinstated into active firing. Neural network models operate by exchanging patterns of spiking activity, and there exists no theory of how computations could unfold solely via transient synaptic changes (Mongillo et al., 2008). Thus, we predicted that, for an activity-silent WM to enter into an information-processing stream, it would first have to be reinstated into an active form.

We evaluated the limits of information processing for active versus activity-silent WM by asking participants to perform a delayed mental rotation task with subjectively seen and unseen stimuli. Our results suggest that this task can be performed even with invisible stimuli, but that such a manipulation of WM involves the reinstatement of consciousness and persistent neural activity, thus suggesting an intrinsic limit to both activity-silent and non-conscious operations.

## Results

We collected behavioral measures in a first set of participants (*n* = 23), then recorded magnetoencephalography (MEG) signals in a second sample (*n* = 30), always employing the same experimental task (**Figure 1**). On each trial, a target square in gray (barely visible target-present trials, 80%) or black ink (target-absent control condition, 20%) was flashed in 1 of 24 possible locations, then masked. Halfway during the ensuing 3 s delay period, a symbolic cue instructed participants to maintain the original target location (no-rotation condition), or to mentally rotate it 120° clockwise or counter-clockwise (rotation condition). Subjects had to comply with these instructions even if they had not seen the target: They were asked to guess the correct final response location if necessary. At the end of a trial, participants rated their subjective visibility of the target using the classical perceptual awareness scale (Ramsøy and Overgaard, 2004), ranging from *1* (no perception whatsoever) to *4* (clearly seen).

**Figure 1.**
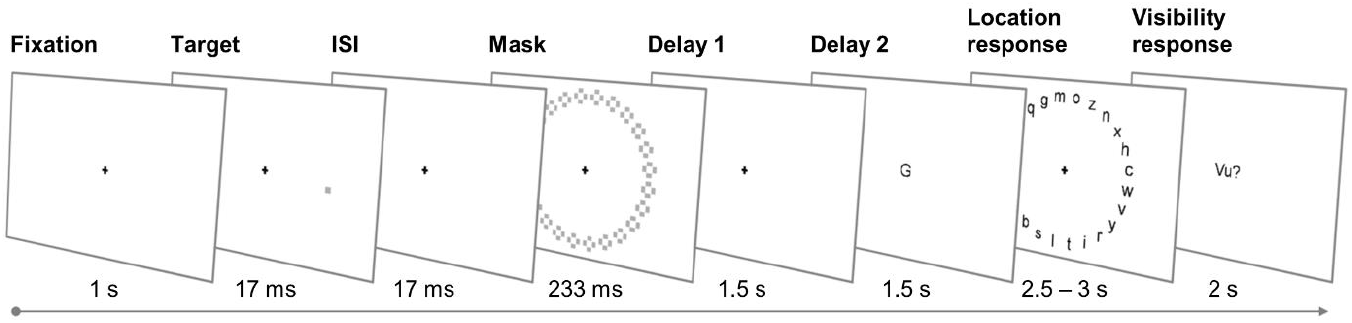
Experimental design. In the behavioral and MEG experiment, participants completed the same spatial delayed-response task. On each trial, a faint target was flashed in 1 out of 24 possible locations and masked. A letter cue presented halfway through a 3 s delay period instructed subjects on the specific task to be performed: (1) Following an equal-sign (« = »), participants were to report the exact location in which the target had appeared. (2) The letter *D* indicated a 120° clockwise, and (3) the letter *G* a 120° counter-clockwise rotation with respect to the target position. At the end of a trial, subjects rated their subjective visibility of the target on a 4-point scale.

### Behavioral evidence for mental rotation of non-conscious stimuli

We first quantified the extent to which subjects could detect, maintain, and manipulate targets in the behavioral experiment. Participants varied their visibility ratings as a function of target presence, reporting the vast majority of target-absent trials as unseen (visibility = *1*; 88.1 ± 3.1%) and ~2/3 of the target-present trials as seen (visibility > *1*; 67.7% ± 3.5%). Target detection *d’* therefore exceeded chance (2.0 ± 0.1; *t*(22) = 13. 2 *p* < .001). Task (no-rotation vs. rotation) did not modulate subjects’ visibility (task x target presence x visibility interaction: *F*(1, 22) = 3.2, *p* = .088), suggesting that participants used the rating scale similarly in both tasks.

Forced-choice localization performance corroborated this interpretation. On seen trials in the no-rotation condition, accuracy was relatively high (65.8 ± 2.5%; chance = 4.17%) and increased monotonically from glimpsed (visibility = *2*) to clearly seen targets (visibility = *4*; all pair-wise comparisons: *p* < .05, except for the comparison between visibility *2* and *3*, where *p* = .296; **Figure 2A, top**). Accuracy remained high on seen rotation trials (30.1 ± 1.9%), albeit, as anticipated, lower than on no-rotation trials (*t*(22) = 12. 3, *p* < .001), and without a clear increase as a function of visibility (all pair-wise comparisons: *p* > .180; **Figure 2A, bottom**). Most crucially, even on the unseen trials, performance was well above chance for the no-rotation and rotation task, irrespective of rotation direction (**Table 1**).

**Figure 2.**
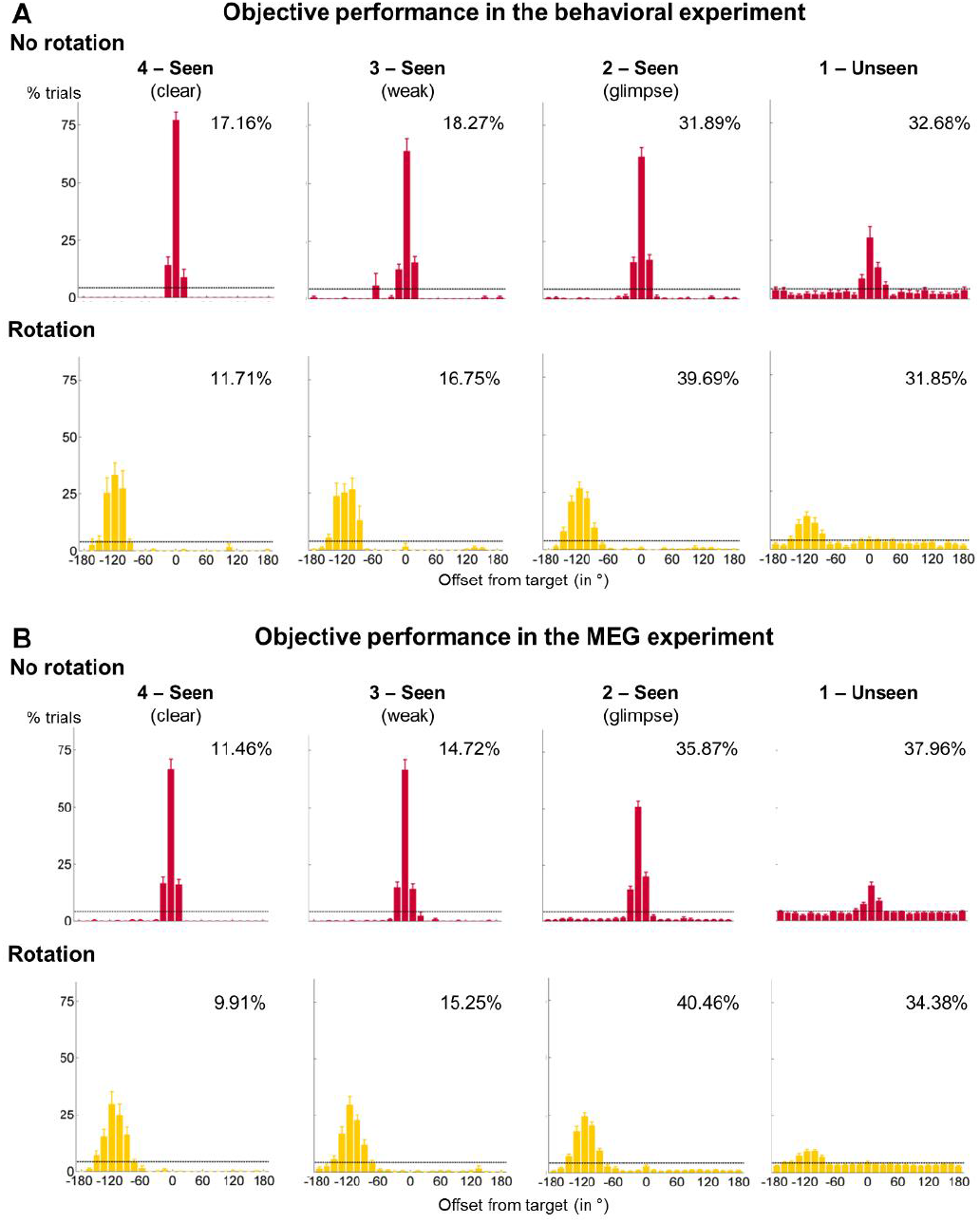
Spatial distributions of forced-choice localization performance in the behavioral (**A**) and MEG (**B**) experiment as a function of task (i.e., no-rotation vs. rotation) and visibility (0° = target location; positive displacement = counter-clockwise offset). The positions at −120° and +120° correspond to the correct locations after clockwise/counter-clockwise rotation. For all analyses and figures, clockwise and counter-clockwise rotations were combined by normalizing all rotation trials into a single rotation condition (i.e., following a counterclockwise rotation, reflecting a position against 0°). Error bars illustrate the standard error of the mean (SEM) across subjects. The horizontal, dotted lines indicate chance at 4.17%. Percentages in the top right corner of each graph show the grand mean proportion of target-present trials from a given visibility category. Due to low number of trials in visibility ratings *2*, *3*, and *4*, we collapsed these ratings into a *seen* category.

**Table 1.** Summary statistics for long-lasting blindsight effect. We display the mean (*M*) and standard error (*SE*) for accuracy on the unseen trials as a function of experiment and rotation condition. *T*-statistic refers to a one-sample test against chance (i.e., 4.17%). Bold numbers indicate significant above-chance localization performance (one-tailed). df = degrees of freedom.

| Experiment | No rotation |  |  | Combined rotation |  |  | Clockwise rotation |  |  | Counter-clockwise rotation |  |  |
| --- | --- | --- | --- | --- | --- | --- | --- | --- | --- | --- | --- | --- |
| | $M$ ($SE$) | $t(df)$ | $p$ | $M$ ($SE$) | $t(df)$ | $p$ | $M$ ($SE$) | $t(df)$ | $p$ | $M$ ($SE$) | $t(df)$ | $p$ |
| Behavior | 26.2% (4.6%) | 4.8 (22) | < .001 | 14.2% (2.1%) | 4.8 (22) | < .001 | 12.5% (2.2%) | 3.7 (22) | < .001 | 16.0% (2.6%) | 4.5 (22) | < .001 |
| MEG | 15.4% (1.5%) | 7.3 (29) | < .001 | 9.2% (1.1%) | 4.3 (29) | < .001 | 9.0% (1.3%) | 3.8 (29) | < .001 | 9.2% (1.3%) | 3.8 (29) | < .001 |

As shown in **Figures 2A** and **3A**, subjects’ responses always surrounded the correct location, yet with greater spread after rotation than no-rotation trials. We separately quantified the rate of approximately correct responding (i.e., correct location ± 30°) and the precision of the spatial representations held in WM (i.e., standard deviation within this tolerance interval; see Methods and Trübutschek et al., 2017). Both task (*F*(1, 22) = 9.9, *p* < .001) and visibility (*F*(1, 22) = 151.1, *p* < .001) affected the rate of correct responding. Participants’ responses fell near the correct location more often in the no-rotation (76.5 ± 2.4%) than in the rotation condition (69.4 ± 2.4%), and when having seen (94.1 ± 1.0%) rather than when not having seen the target square (51.9 ± 3.8%). These factors did not interact (*F*(1, 22) = 0.2, *p* = .657; **Figure 3A, top inset**), indicating that decrements in performance following a mental rotation were comparable across seen and unseen targets.

**Figure 3.**
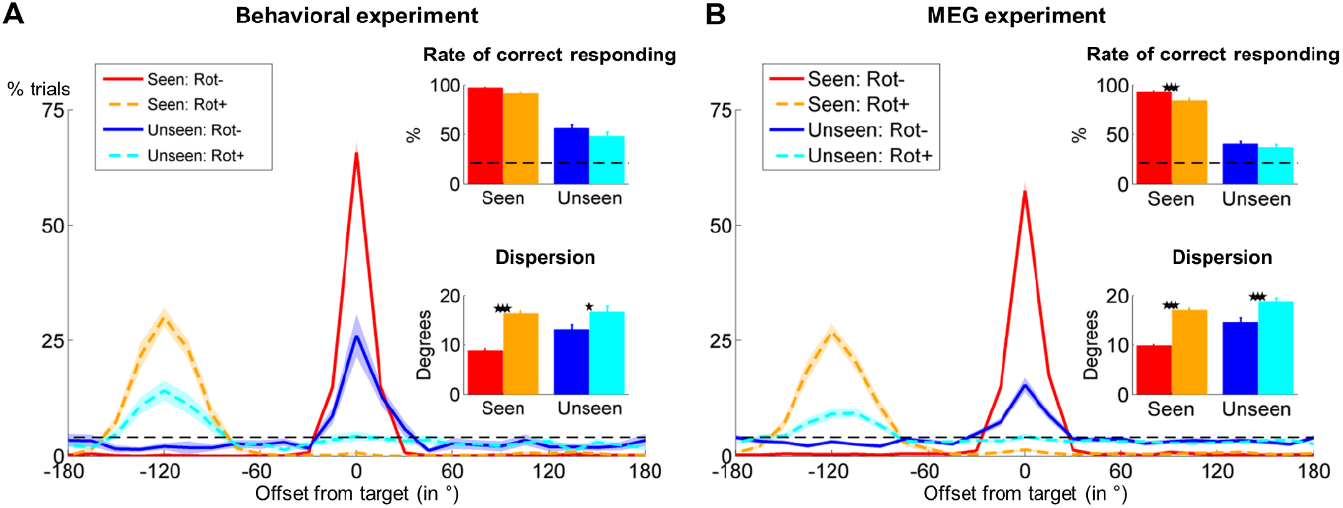
Behavioral evidence for manipulation of non-conscious information in the behavioral (**A**) and MEG (**B**) experiment. Panels depict distributions of participants’ localization responses with respect to the target location (0°; positive displacement = counter-clockwise offset) as a function of task (no rotation = solid line, rotation = dotted line) and visibility (seen = warm colors, unseen = cool colors). Insets show the rate of correct responding (proportion of trials within ± 2 positions of correct response location; top) and the precision of working-memory representations in all participants with sufficient blindsight (bottom). Horizontal dotted lines index chance at 4.17% (for single locations) and 20.83% (for the region of correct responding) respectively. Shaded area and error bars represent the standard error of the mean (SEM) across subjects. *p < .05, ** < .01, and *** p < .001 in a paired samples *t*-test.

Analysis of precision reinforced this conclusion: Out of 23 subjects, 19 displayed above-chance blindsight across both rotation directions (chance = 20.83%; *p* < .05 in a *χ*^2^-test) and were thus included here. Task (*F*(1, 18) = 34.9, *p* < .001) and visibility (*F*(1, 18) = 10.3, *p* = .005) again influenced localization performance, but this time also interacted (*F*(1, 18) = 8.9, *p* = .008). Rotating the target location decreased the precision of participants’ responses for seen (*t*(18) = −11.9, *p* < .001) and unseen targets (*t*(18) = −2.3, *p* = .031), but this reduction was stronger for seen than unseen trials (*t*(18) = −3.0, *p* = .008; **Figure 3A**, **bottom inset**). Again, there was therefore no observable detriment to rotating an unseen location.

We replicated these observations in the MEG experiment. Subjects employed the visibility scale meaningfully, rating target-present trials primarily as seen (64.6 ± 3.2%) and target-absent trials as unseen (83.6 ± 2.5%; detection *d’:* 1.7 ± 0.1, *t*(29) = 14.2, *p* < .001) in both tasks (task x target presence x visibility interaction: *F*(1, 29) = 2.1, *p* = .159). Localization accuracy for seen targets was modestly high in the no-rotation condition (57.5 ± 2.2%; **Figure 2B, top**) and reduced following a mental rotation (27.1 ± 1.6%, *t*(29) = 14.3, *p* < .001; **Figure 2B, bottom**). Again, we observed a long-lasting blindsight effect in both tasks and for all rotation directions (**Table 1**). Task and visibility influenced the rate of correct responding (main and interaction effects: all *F*s(1, 29) > 4.8, all *p*s < .036) and precision (n = 27; main and interaction effects: all *F*s(1, 26) > 8.3, all *p*s < .008). Mental rotation decreased participants’ performance on seen (*t*(29) = 5.0, *p* < .001), but not on unseen trials (*t*(29) = 1.8, *p* = .090; **Figure 3B, top inset**), and also reduced precision more following a rotation with seen (*t*(26) = −15.9, *p* < .001) than unseen targets (*t*(26) = −3.9, *p* < .001; **Figure 3B, bottom inset**).

These findings show that, even when failing to perceive the target, subjects succeeded in manipulating it. However, there exist at least three possible explanations for this long-lasting blindsight effect. First, it may have been the product of a genuine non-conscious manipulation. Second, it may have resulted from a fraction of *seen* trials miscategorized as *unseen*, yet still yielding correct performance; this interpretation, although rejected in our previous experiment without rotation (Trübutschek et al., 2017), needs to be re-examined here. Third, subjects may have recovered the information from non-conscious WM around the time of the cue, transformed it into a conscious, active representation (forced-choice retrieval) and thereafter consciously manipulated this early guess. To resolve these possibilities, we turned to our MEG data, focusing on five a-priori time windows: early brain responses (0.1 – 0.3 s), the P3b time window previously shown to be critical for conscious perception (0.3 – 0.6 s), the delay period before (0.6 – 1.76 s) and after (1.76 – 3.26 s) the rotation cue, and the response period (3.26 – 3.5 s).

### Long-lasting blindsight does not arise from miscategorization of seen trials

Above-chance objective performance for unseen targets could have resulted from the erroneous mislabeling of some seen targets as unseen. If this were the case, the unseen correct trials should display the same neural signatures of conscious processing as seen trials (Trübutschek et al., 2017). There should be an amplification of brain activity during the P3b time window, and a classifier trained to distinguish accuracy on the unseen trials should resemble a standard visibility decoder (i.e., seen vs. unseen). By contrast, the classification of seen versus unseen correct trials should produce a different pattern of results or fail entirely.

To evaluate this alternative miscategorization hypothesis, we first characterized univariate neural markers tied to conscious perception. Contrasting brain activity on seen and unseen trials revealed typical signatures of conscious processing (Gaillard et al., 2009; Sergent et al., 2005; Trübutschek et al., 2017). Seen targets elicited a strong positive response between ~300 and 600 ms in right-lateralized centro-parietal sensors, corresponding to activations in occipital, temporal, parietal and dorsolateral prefrontal brain areas (*p*_clust_ = .011; **Figure 4A**). Moreover, brain activity was amplified during the P3b time window (i.e., ~292 and 576 ms; *p*_uncorrected_ < .05), though further differences with unseen targets also persisted between ~964 and 1320 ms (*p*_uncorrected_ < .05; **Figure 4B**). Importantly, task did not modulate these brain responses (task x visibility interaction: *p*_clust_ > .280).

**Figure 4.**
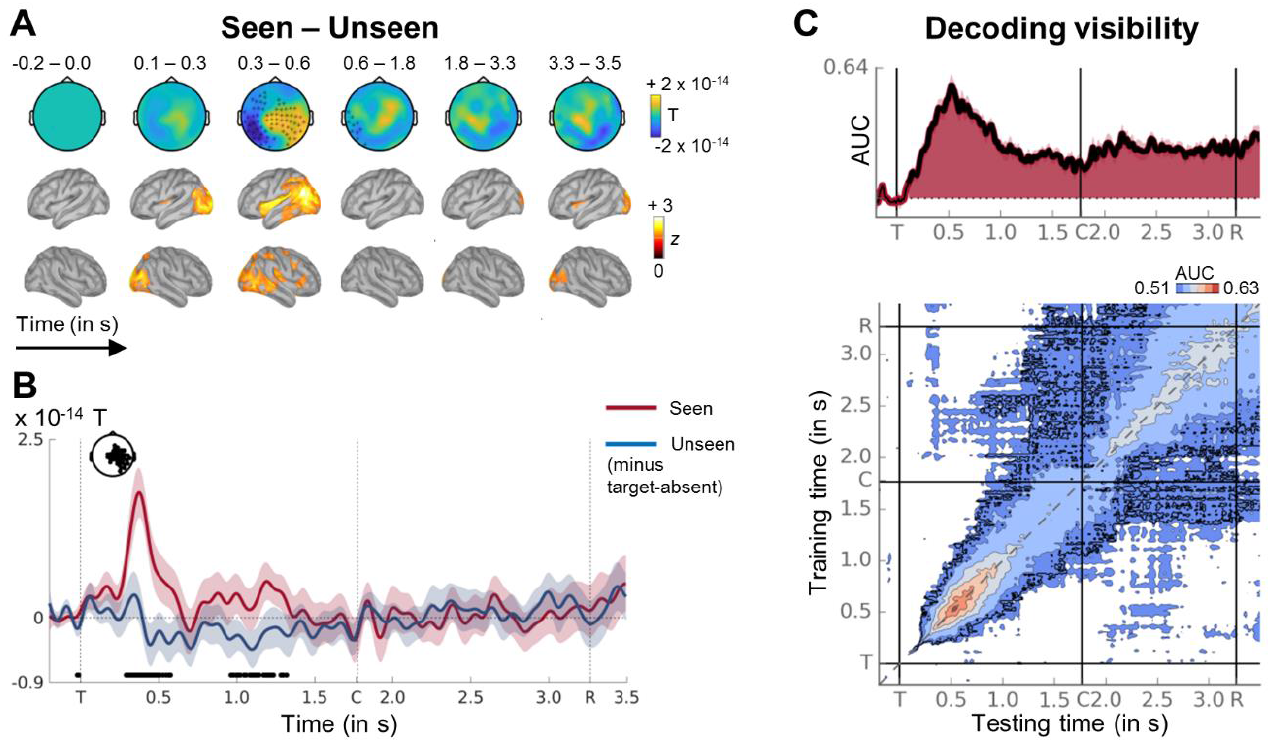
Typical neural signatures and dynamics of conscious processing for seen targets. **(A)** Sequence of brain activations (−0.2 – 3.5 s) evoked by seen targets in both tasks in sensor (top) and source space (bottom). Each topography depicts the difference in amplitude between seen and unseen trials averaged over the time window shown (magnetometers only). Sources reflect z-scores of absolute difference with respect to a pre-stimulus baseline. Black asterisks indicate sensors showing a significant difference between seen and unseen trials at any point during the respective time window as assessed by a Monte-Carlo permutation test. **(B)** Average time courses (−0.2 – 3.5 s) of seen (red) and unseen (blue) trials in that subset of magnetometers having shown a significant effect in (A). Shaded area illustrates standard error of the mean (SEM) across subjects. Significant differences between conditions are depicted with thick black line (two-tailed Wilcoxon signed-rank test, uncorrected). Vertical dotted lines index onset of the target (T), symbolic cue (C), and response (R) screens. For display purposes only, data were lowpass-filtered at 8 Hz. **(C)** (Top) Average time course of diagonal decoding of visibility (i.e., seen vs. unseen). Thick black line and shaded area denotes above-chance decoding as assessed by a one-tailed cluster-based permutation analysis. Horizontal, dotted line represents chance level at 50%. (Bottom) Temporal generalization matrix of the same visibility decoder. Each horizontal row in the matrix corresponds to an estimator trained at time *t* and tested on all other time points *t*’. The diagonal gray line demarks classifiers trained and tested on the same time points (i.e., the diagonal estimator shown on top). Thick black outline indexes above-chance decoding as evaluated by a two-tailed cluster-based permutation test. In both plots, vertical lines mark onset of the target (T), symbolic cue (C), and response (R) screens. For display purposes, data were smoothed with a moving average of 5 samples (i.e., 40 ms). AUC = area under the curve. See also Figures S1, S2, and S3.

When contrasting the unseen correct with the unseen incorrect epochs, we observed no evidence for a miscategorization. No significant differences emerged (*p*_clust_ > .221) and there was no sign of any amplification of brain activity (**Supplementary Figure 1A**), even when considering the time courses in channels most sensitive to divergences in amplitude for seen and unseen targets (**Supplementary Figure 1B**). Bayesian statistics provided substantial evidence in favor of the null hypothesis (i.e., no difference in MEG amplitude between unseen correct and incorrect trials) for all time windows (all Bayes’ Factors < 0.38).

Because chance corresponded to 20.83% (i.e., 5/24 positions), a non-negligible portion of the unseen correct trials might have resulted from guessing, thus potentially obscuring differences between unseen correct and incorrect epochs. To address this possibility, we next estimated neural activity for unseen correct epochs while accounting for chance-responding (cf. Lamy et al., 2009, footnote 2). If these chance-free unseen correct trials resulted from a miscategorization of seen epochs, we should now observe clear signatures of conscious processing. This was not the case. Chance-free brain activity was still indistinguishable from the one on unseen incorrect and unseen correct trials (whole-brain: all *p*_clust_ > .252; critical time courses: all Bayes’ Factors < 0.76). Moreover, it remained strikingly different from a synthetic waveform, derived by proportionally mixing the signals from seen and unseen incorrect trials (as would be expected under the miscategorization hypothesis; **Supplementary Figure 1B**). Those findings allow us to reject the hypothesis of a miscategorization of some seen trials as unseen.

Decoding analyses refined this conclusion. Training a linear multivariate pattern classifier to discriminate seen from unseen trials resulted in above-chance diagonal decoding from ~120 ms to the end of the epoch (all *p*_clust_ < .05; **Figure 4C, top**), quickly peaking at ~528 ms, then first slowly decaying until the cue before being sustained throughout the remainder of the trial (time bins: AUCs > 0.54, *p*s_corr_ < .005). The temporal generalization of each estimator trained at a specific time to all other time points confirmed this picture (**Figure 4C, bottom**): Visibility decoding was primarily confined to a thick diagonal, indicating that conscious perception was associated with a dynamically evolving chain of metastable patterns of brain activity (King and Dehaene, 2014). Similar findings emerged when training and testing a visibility classifier separately in the no-rotation and rotation condition, or when generalizing from one task to the other (**Supplementary Figure 2**). Multivariate neural signatures of conscious perception were thus stable across experimental tasks and in line with previous observations (Marti et al., 2015; Salti et al., 2015; Trübutschek et al., 2017).

Crucially, we found no discernable pattern when classifying unseen correct versus unseen incorrect trials (all *p*_clust_ > .05; time bins: AUCs < 0.51, *p*s_corr_ > .05; Bayes’ Factors < 0.28; **Supplementary Figure 1C**). However, training a classifier to distinguish the seen from the unseen correct epochs resulted in a similar, albeit weaker, decoding time course and generalization matrix as when directly training on all unseen or even just the unseen incorrect trials (time bins: AUCs > 0.52, all *p*s_corr_ < .05; Bayes’ Factors > 2.07; **Supplementary Figure 3**). As such, this pattern of results is exactly opposite to what one would have expected in the case of a miscategorization. These findings persisted even when including only those subjects with sufficient blindsight (*n* = 27). This replication of our previous work (Trübutschek et al., 2017) thus rules out a miscategorization of unseen correct trials as an alternative explanation for the long-lasting blindsight effect. Instead, it indicates that information was genuinely encoded in non-conscious WM.

### Long-lasting blindsight effect results from active, conscious rotation

What process allowed participants to perform a mental rotation on unseen trials? Was it the result of a genuine non-conscious manipulation? Or did subjects perform a conscious manipulation by first reinstating an active representation of the estimated target position around the time of the rotation cue and then rotating this conscious guess? Disambiguating between these alternatives requires the identification of a neural marker of active, conscious processing. Prior work has pointed towards a rhythmic signal – a suppression of power in the alpha (8 – 12 Hz) and low (13 – 20 Hz) as well as high beta frequency bands (20 – 27 Hz) – as a reflection of such a cognitive state (Backer et al., 2015; Meyniel and Pessiglione, 2014; Trübutschek et al.,İ2017).

Across all trials, we indeed observed a prominent desynchronization in alpha/beta frequencies over an extensive set of central sensors, emanating primarily from parietal brain sources (**Figure 5A**). Cluster-based permutation analyses revealed reliable differences in brain responses in a slightly larger set of channels between seen targets and all other experimental conditions exclusively prior to the presentation of the rotation cue. Power decreased more strongly on seen than on unseen trials between ~580 and 1320 ms in the alpha (*p*_clust_ = .032), and between ~460 and 1300 ms in the low beta band (*p*_clust_ = .046; **Figure 5B, top**). Similarly, pre-cue desynchronizations were more pronounced for seen than for target-absent epochs in the low (*p*_clust_ = .015) and high beta bands (*p*_clust_ = .030) between ~280 and 940 and ~820 and 2000 ms. There were no discernable differences in the power profiles between (1) unseen and target-absent trials (all *p*_clust_ > .250) and (2) unseen correct and incorrect epochs (all *p*_clust_ > .280; **Figure 5B, bottom**). Desynchronization of alpha/beta power may therefore serve as a signature of conscious processing in the current task.

**Figure 5.**
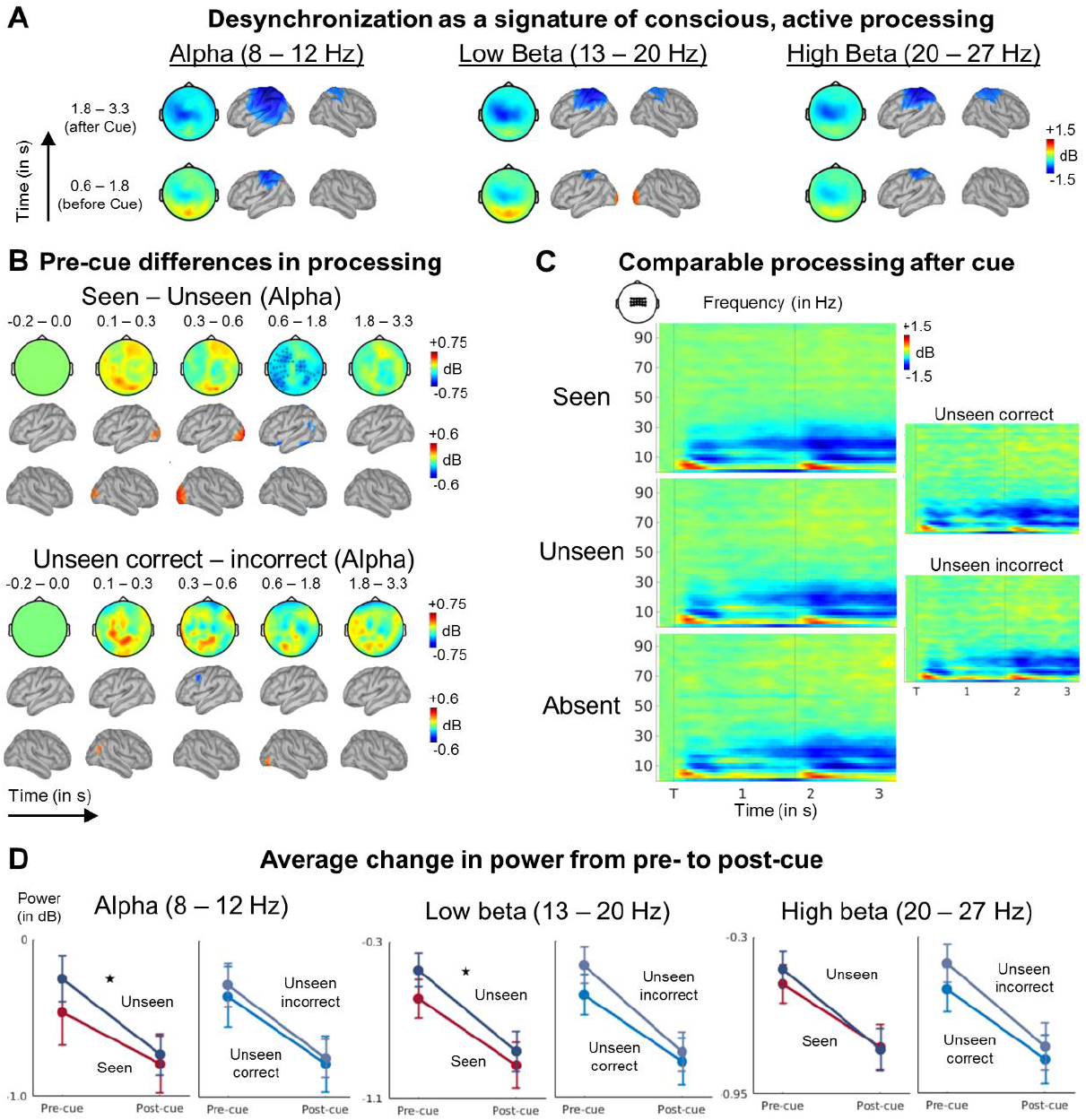
Time-frequency markers of conscious processing emerge around the time of the rotation cue on the unseen trials. (**A**) Average pre-cue (0.6 – 1.8 s; bottom) and post-cue (1.8 – 3.3 s; top) desynchronization in the alpha (8 – 12 Hz; left), low beta (13 – 20 Hz; middle), and high beta (20 – 27 Hz; right) frequency bands in magnetometers and source space (in dB; relative to pre-stimulus baseline). (**B**) (Top) Alpha band activity (8 – 12 Hz) related to consciously perceiving the target square (i.e., seen vs. unseen) is shown in magnetometers and source space (in dB; relative to pre-stimulus baseline). Black asterisks denote cluster of sensors displaying a significant difference at any point in time during the respective time window (as evaluated by a Monte-Carlo permutation test. (Bottom) Same as on top, but for the contrast between unseen correct and unseen incorrect trials. (**C**) Average time-frequency power relative to baseline as a function of visibility and target presence in a subset of central magnetometers. Horizontal lines demark onset of target (T) and cue presentation. (**D**) Plots depict average pre-cue and post-cue power in the same group of sensors as in (C) as a function of frequency (i.e., alpha, low beta, and high beta) and visibility (i.e., seen, unseen, unseen correct and unseen incorrect). Error bars represent standard error of the mean (SEM) across subjects. Asterisks denote significant interaction in a repeated-measures ANOVA at *p* < .05. See also Figure S4.

Using this marker, we are now in a position to evaluate the remaining alternatives. If the long-lasting blindsight effect resulted from a genuine, non-conscious rotation, on seen trials, we should observe a sustained desynchronization in the alpha and beta bands throughout the entire epoch, while no (or at least significantly weaker) power decreases should be associated with unseen and target-absent epochs. By contrast, if participants consciously rotated a guess, neural signatures of conscious processing should be highly similar across all experimental conditions after the cue. Differences in desynchronization between seen and unseen/target-absent trials should only exist during the pre-cue phase.

Our results support the latter hypothesis (**Figure 5C**). Following an initial divergence during the early pre-cue maintenance phase (**Supplementary Figure 4A-C**), differences in spectral profiles between seen, unseen, and target-absent trials vanished by ~1 s. All epochs were characterized by a prominent, sustained desynchronization in the alpha, low and high beta frequencies. This suppression in power varied as a function of subjective visibility (i.e., seen vs. unseen) and time (i.e., pre-cue vs. post-cue delay). It was much more pronounced during the post-cue than the pre-cue maintenance period (i.e., main effect of time: all *F*s > 18.6, all *p*s < .001). Crucially, this difference between pre- and post-cue power was also larger for unseen than for seen targets in the alpha and low beta bands (visibility x time interaction: all *F*s > 4.01, all *p*s <= .05), and marginally so in the high beta band (visibility x time interaction: *F*(1, 29) = 2.95, *p* = .097; **Figure 5D**). No such interaction emerged when contrasting the unseen correct with the unseen incorrect trials (i.e., visibility x time interaction: all *F*s < 2.83, all *p*s > .103; **Figure 5D**), as these conditions displayed largely similar power profiles throughout the entire epoch (**Supplementary Figure 4D-F**).

We thus observed a reliable distinction between seen and unseen brain states only during the maintenance period preceding the execution of the experimental task up until at least 1 s. Seen targets were accompanied by a significantly larger desynchronization in the alpha and low as well as high beta frequencies. These differences vanished entirely by the time the symbolic rotation cue was presented. The mental rotation task appeared to be solved by reinstating a conscious estimate of target location.

### The location of unseen targets can only be tracked transiently

To further test this conclusion, we used multivariate decoding to track neural activity underlying the encoding, maintenance, manipulation and retrieval of seen and unseen target locations. We first trained a multivariate regression model to predict target angle from participants’ brain activity separately for each point in time. In order to maximize statistical power and increase our ability to detect small effects, we fitted the estimator while collapsing target-present trials across rotation and visibility conditions. We then evaluated model performance on left-out subsets of epochs (see Methods for details). Note that, unless explicitly stated, none of the findings changed qualitatively when testing separately on the rotation and no-rotation task (**Supplementary Figure 5**).

Starting at ~80 ms, estimator performance for seen targets steadily rose until ~264 ms and then slowly decayed towards chance at ~1.46 s (**Figure 6A**). Following the rotation cue, a rebound of position-selective activity was observed and was then fairly sustained for the remainder of the trial, with a short gap between ~2.70 and 3.10 s right before the onset of the response screen (*p*_clust_ < .05; time bins: *W*s > 417.0, *p*s_corr_ < .005, Bayes’ Factors > 77.93). Thus, in line with previous findings (Trübutschek et al., 2017), seen targets were initially encoded via active neural firing. Then, this representation decayed and was reactivated throughout most of the post-cue delay period.

**Figure 6.**
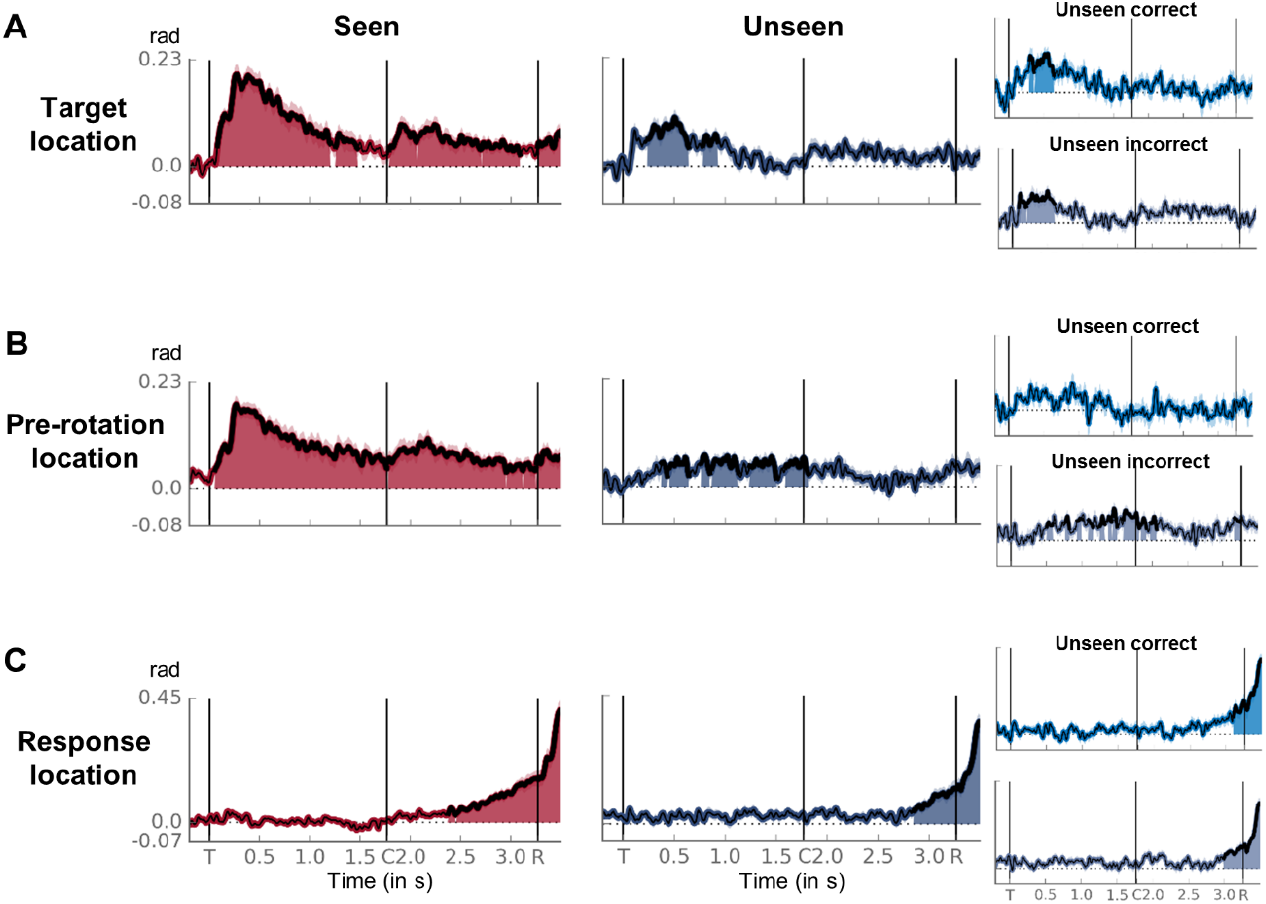
Tracking a mental rotation on seen and unseen trials. (**A**) Time courses of average decoding of target location on seen (red), unseen (dark blue), unseen correct (light blue) and unseen incorrect (blue) trials. Thick lines and shaded areas represent above-chance performance as assessed by a one-tailed cluster-based permutation test. Horizontal dotted lines index chance. Event markers denote the onset of the target (T), cue (C), and response (R) screens. For illustration purposes, data were smoothed with a moving average of 5 samples (i.e., 40 ms). (B) Same as in (A), but for pre-rotation location. (**C**) Same as in (A), but for response location. See also Figure S5.

A different picture emerged for unseen targets. While target location was again encoded and actively stored during the early part of the epoch, this representation was weaker than the one for seen targets (paired-samples Wilcoxon signed rank test: pre-cue time bins: *W*s > 370.0, *p*s_corr_ < .02, Bayes’ Factors > 3.42) and decayed much more quickly, vanishing entirely by ~920 ms (*p*_clust_ < .05; pre-cue time bins: *W*s > 351.0, *p*s_corr_ < .035, Bayes’ Factors > 7.34). During the post-cue delay period, although we found no evidence in favor of an actively coded representation of target location when considering the decoding time course itself (*p*_clust_ > .05; **Figure 6A**), the estimator’s performance over the entire time window remained above chance (rads = 0.03 ± 0.01, *W* = 355.0, *p*_corr_ = .025, Bayes Factor 6.41) and at comparable levels as on seen trials (*W* = 315.0, *p*_corr_ = .460, Bayes’ Factor = 0.86). A more fine-grained analysis with a moving average of 100 ms revealed that this effect was driven primarily by the initial phase of the delay, up to ~2.6 s. We observed no modulation of this pattern of findings by accuracy (time bins: *W*s < 279.0, all *p*s_corr_ > .950, Bayes’ Factors < 0.41; **Figure 6A, insets**).

Overall then, a mixture of two different mechanisms seems to have supported the initial, pre-cue storage of seen and unseen target locations. Whereas seen targets were maintained with persistent albeit decaying, neural activity, unseen targets elicited weaker position-related activity that also quickly decayed to baseline-level. During the post-cue phase, once participants either actively maintained or manipulated the contents of their WM, the representation of seen targets was reactivated and sustained for the remainder of the epoch. Unseen targets may also have benefitted from a short-lived revival, but this effect was weak and the associated decoding time course much less compelling than the one for seen trials.

### An estimate of the location of unseen targets is reinstated prior to the rotation cue

Localization responses on unseen trials did not always follow the actual target position. On more than half of the unseen trials (62.0 ± 2.8%), subjects chose an incorrect location. What determined participants’ final response on those trials? According to the activity-silent account of WM, around the time of mental rotation, subjects should have attempted to reinstate an active neural representation of the target, albeit with occasional location errors, and then rotated this guess. To evaluate this prediction, we set out to track the neural representation of participants’ location estimates throughout the task. Around the time of the rotation cue, brain signals should contain a decodable representation of the “pre-rotation location”, i.e. the spatial location that, given the subjects’ response, would have been the location retrieved and then rotated. On no-rotation trials, this location coincided with response location, whereas on rotation trials, it corresponded to the position of participants’ response rotated 120° in the direction opposite to what the rotation cue had instructed. Detecting the presence of such a pre-rotation representation on unseen rotation trials would support the results of our time-frequency analyses and the hypothesis that, around the time of the cue, subjects attempted to recover a conscious representation of the target (sometimes an erroneous one) and then consciously rotated this guess. If, however, unseen performance was based on an active manipulation of activity-silent WM, then such decoding should fail.

On seen trials, decoding the pre-rotation location was possible, with a time course strikingly similar to the one for the true position of the target (**Figure 6B**). From ~56 ms onwards, the pre-rotation location was coded in activity-based brain states (*p*_clust_ < .05; time bins: *W*s > 408.0, *p*s_corr_ < .005, Bayes’ Factors > 517.26), first peaking at ~ 264 ms (rad = 0.18 ± 0.02) and then slowly decaying before being revived by the rotation cue and sustained for the remainder of the epoch.

Crucially, pre-rotation location could also be decoded on unseen trials. Shortly after the presentation of the target, the estimator’s performance began to rise and first exceeded chance at ~376 ms (rad = 0.052 ± 0.015). Decoding persisted until ~1.8 s (*p*_clust_ < .05; P3b time window and pre-cue delay: *W*s > 382, *p*s_corr_ < .005, Bayes’ Factors > 78.83), though estimator performance itself did not drop until ~ 2.5 s. Indeed, a follow-up analysis with narrower 100-ms time windows suggested that the pre-rotation location may have been maintained until ~ 2.2 s *(p* < .05, uncorrected). There was again no evidence for a modulation of this pattern as a function of accuracy (time bins: *W*s > 120.0, *p*s_corr_ > .600, Bayes’ Factors < 1.44; **Figure 6B, insets**).

As predicted, while the representation of the pre-rotation location was stronger for seen than for unseen targets during the early part of the epoch (early and P3b time window: *W*s > 450.0, *p*s_corr_ < .005, Bayes’ Factors > 124,688.30), this difference started to diminish during the pre-cue maintenance phase (*W* = 347.0, *p*_corr_ = .085, Bayes’ Factor = 1.76) and vanished entirely by the last second before the rotation cue (moving average of 100 ms: *W*s < 359.0, *p*s_corr_ > .05, Bayes’ Factor < 1.32). Participants’ location estimates were therefore similarly represented on both seen and unseen trials during the last part of the pre-cue maintenance period: Even on unseen trials, the material rotated was an active, conscious guess of a target location.

### An active representation of target location is mentally rotated in WM

We last trained and tested a multivariate regression model to decode response location. On seen trials, response location emerged reliably only in the second half of the post-cue delay period (**Figure 6C**). Starting at ~2.38 s, decoding performance gradually built up until its peak at the very end of the epoch (*p*_clust_ < .05; post-cue time bins: *W*s > 440.0, *p*s_corr_ < .005, Bayes’ Factors > 21,997.68). There was substantial temporal overlap between the decoding of the target/pre-rotation location and the response position: As the former started to decay around ~2.5 s, the latter slowly began to pick up.

**Figure 7** further shows the probability density distributions for decoded target and response locations. On seen trials, prior to the rotation cue, decoder estimates for target angle were strongly concentrated around the actual target location, irrespective of rotation condition and direction (resultant vector lengths > .41; Rayleigh tests for non-uniformity: zs > 5.09, *p*s < .005; non-parametric multi-sample test for equal medians: *p*s > .302). This picture changed following the rotation cue. While angle estimates on no-rotation trials continued to stay fairly centered on the original target location (resultant vector lengths > .37; Rayleigh test: z > 4.01, *p* < .017), their counterparts for clock- and counter-clockwise rotations began to shift towards the respective correct response positions (response period: clockwise rotation: *M*_circ_ = 37.3°; resultant vector length = .49; one-sample test against a mean direction of 0°: *p* < .05; counterclockwise rotation: *M*_circ_ = 95.6°; resultant vector length = .31; one-sample test against a mean direction of 0°: *p* < .05). During the response period, all three distributions were characterized by a different center of mass (non-parametric multi-sample test for equal medians: *p*s < .05), located in close proximity to the expected final position. Depending on the direction of the rotation, the representation of the original target location was progressively transformed into a representation of the response position. On average, then, a mental rotation following seen targets was reflected by an active transition period, during which the stimulus code was progressively replaced by the response code. Note however that, while such a smooth transition was visible in the mean, we cannot determine here whether continuous or discrete transitions occurred on individual trials (Latimer et al., 2015).

**Figure 7.**
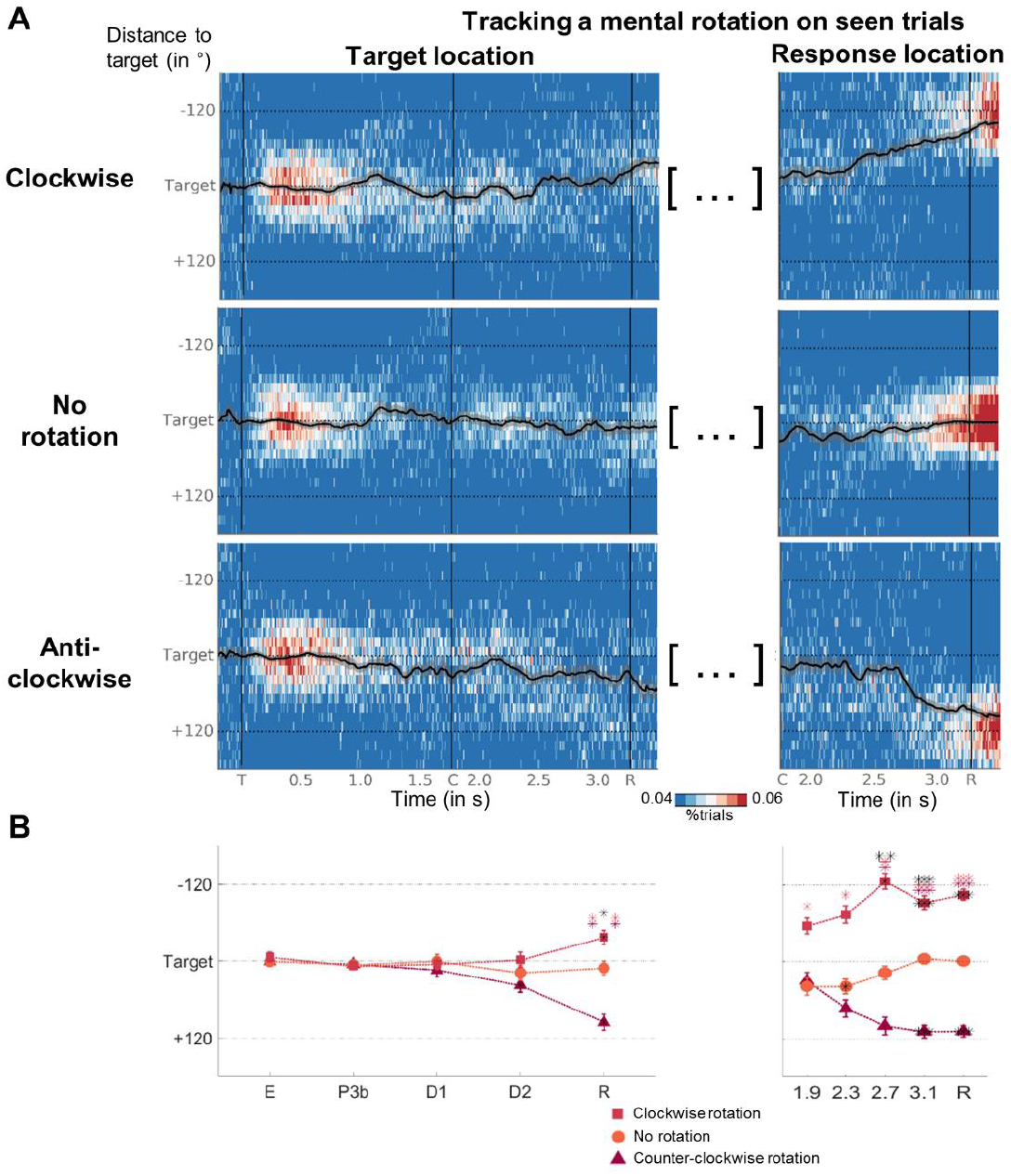
Tracking a mental rotation on seen trials. **(A)** (Left) Time courses of probability density distributions of the angular distance between the estimates of a decoder trained with target angle and actual target location are shown as a function of rotation condition. For display purposes, data were smoothed with a moving average of 12 samples (i.e., 96 ms). Overlaid black line illustrates the evolution of the circular mean of the individual distributions (also smoothed). Shaded area reflects circular standard variation across subjects. Vertical event markers denote the onset of the target (T), cue (C), and response (R) screens, horizontal markers index correct response positions after rotation. (Right) Same as in the left panels, except for angular distance between the estimates of a decoder trained with response angle and actual target location. **(B)** Circular means of the above distributions as a function of rotation condition and time bin (i.e., E = 100 – 300 ms, P3b = 300 – 600 ms, D1 = 0.6 – 1.76 s, D2 = 1.76 – 3.26 s, R = 3.26 – 3.5 s). Error bars reflect circular standard deviation. Asterisks inside markers denote significant deviation from mean direction of 0 (as assessed by a circular equivalent of a one-sample t-test), asterisks on top significant differences in median direction between conditions (as assessed by a circular equivalent to the Kruskal-Wallis test; black = clockwise vs. counter-clockwise; red = clockwise vs. no rotation; violet = counter-clockwise vs. no rotation). * *p* < .05, ** *p* < .01, *** *p* < .001. See also Figure S6.

We next considered the unseen trials. If subjects similarly performed a conscious rotation of (an estimate of) unseen locations, then one would predict the response estimator to perform comparably on seen and unseen targets. This was indeed the case (**Figure 6C**). Decoding response location on unseen trials yielded consistent above-chance performance from ~2.84 s onwards (*p*_clust_ < .05; post-cue time bins: *W*s > 410.0, *p*s_corr_ < .005, Bayes’ Factors > 594.74), again beginning to rise around the same time as the model for the pre-rotation location had faded (cf. time courses in **Figure 6B** and **C**). As would be expected if the same underlying process were responsible for the generation of responses across all experimental conditions, we observed no differences as a function of accuracy (time bins: *W*s < 314.0, *p*s_corr_ > .480, Bayes’ Factors < 0.81) or visibility (time bins: *W*s < 334.0, *p*s_corr_ > .600, Bayes’ Factors < 2.45). Prerotation and response locations could also be tracked on unseen trials, albeit, as expected, with reduced accuracy (**Supplementary Figure 6**). The transformation from one representation into another therefore appeared to have been comparable for seen and unseen targets, in both cases relying on decodable activity patterns rather than on activity-silent brain states.

## Discussion

Recent work has challenged classical views of WM as a purely conscious process based on persistent neural firing. Instead, information may also be stored in non-conscious, activity-silent WM, without any accompanying neural activity, via slowly decaying changes in synaptic weights (Mongillo et al., 2008; Rose et al., 2016; Stokes, 2015; Trübutschek et al., 2017; Wolff et al., 2015, 2017), and in the complete absence of subjective awareness (Bergström and Eriksson, 2017; Soto et al., 2011; Trübutschek et al., 2017). So far however, only the short-term maintenance of information has been explored, while its transformation, a key feature of WM, has been ignored.

Here, we show that, whether or not information was consciously perceived, manipulating it was associated with a prior reinstatement of an active neural representation, accompanied by signatures of a conscious state. These findings question the term non-conscious *working* memory, and suggest that WM manipulation requires a conversion from activity-silent to active WM.

### Manipulation as a limit for non-conscious, silent processes

It has proven notoriously difficult to put clear upper bounds on the depth of non-conscious processing. Non-conscious signals tend to affect a wide range of behaviors and trigger activity in many different brain areas, including the prefrontal cortex (van Gaal et al., 2010; Naccache and Dehaene, 2001; Nakamura et al., 2018; van Vugt et al., 2018). Recent work on non-conscious WM has even called into question some of the most basic assumptions regarding the nature of non-conscious processes, suggesting that non-conscious signals may be maintained much longer than previously thought (Bergström and Eriksson, 2017; King et al., 2016; Soto et al., 2011; Trübutschek et al., 2017).

Our behavioral results, superficially, support this conclusion, as they provide evidence for a non-conscious process of mental rotation. On unseen trials, subjects reported the correct response position much better than chance after several seconds, irrespective of whether they just had to maintain the original target location or rotate its position. We replicated this long-lasting blindsight effect in two independent experiments and, as such, seemingly expanded the range of possible non-conscious WM processes to include manipulation of information (Bergström and Eriksson, 2015; Bona et al., 2013; Soto et al., 2011; Trübutschek et al., 2017).

Our neural data further indicated that subjective visibility reports were genuine. Prior to the rotation cue, we observed typical markers of conscious, active processing almost exclusively for seen targets. Brain activity was amplified during the P3b time window (Gaillard et al., 2009; Sergent et al., 2005), and participants’ visibility (i.e., seen vs. unseen) was decodable with high accuracy (King et al., 2016; Salti et al., 2015; Trübutschek et al., 2017). Moreover, there was a sustained desynchronization of alpha/beta frequency, which became even more pronounced after the rotation cue, thereby coinciding with the most demanding phase of our task (Pessiglione et al., 2007; Trübutschek et al., 2017; Wyart and Tallon-Baudry, 2009). By contrast, for unseen targets, signatures of conscious processing were entirely absent or markedly reduced in comparison to the ones on seen trials early during the epoch. There was neither an ignition of brain activity during the P3b time window, nor a comparably strong alpha/beta desynchronization. These findings, in line with our previous work (Trübutschek et al., 2017), show that “unseen” trials were genuine and did not correspond to a subset of miscategorized seen trials.

Those neural signatures, however, changed drastically around the time of the mental rotation cue, suggesting that an estimate of target location was reactivated and regained consciousness. Slightly before the rotation cue, around ~1 s, alpha/beta power decreased for unseen targets, reaching similar levels as on seen trials during the post-cue maintenance period. Starting at more or less the same time (i.e., around ~500 ms), a decodable representation of the pre-rotation location emerged. Participants therefore seem to have estimated and reinstated an active representation of target location in anticipation of the upcoming rotation task. On unseen trials, the weak activity-silent representation of the target may have competed against other ongoing noise fluctuations in the brain, resulting in a mixture of trials where decision was solely based on stochastic events (Vul et al., 2009) and others biased towards the correct target location. Variability across trials and participants as well as the temporal smoothing inherent to time-frequency analyses precludes a definitive determination of the exact onset of the pronounced and sustained alpha/beta desynchronization on unseen trials, but the results indicate that this transition already occurred shortly before the presentation of the symbolic rotation cue.

In conjunction with previous work (Bergström and Eriksson, 2017; Soto et al., 2011; Trübutschek et al., 2017), these findings thus highlight the limits of non-conscious WM. While information may be temporarily stored non-consciously, manipulating items is associated with a reinstatement of an active conscious representation. Our results may thus help to circumscribe the boundaries of non-conscious processing. Consciousness has been theorized and empirically demonstrated to be a necessary prerequisite for the execution of serial tasks, such as the chaining of mental operations (Dehaene, 2001; Sackur and Dehaene, 2009). We here observed that such chaining may remain possible even if the initial input was not represented consciously, but only inasmuch as subjects willfully operate on previously non-conscious information by forcing it into an active state before routing it to a conscious processor. Future research might expand on this work and attempt to more strongly encourage the reliance on non-conscious processing by, for instance, rendering the task cues subliminal.

### The complementarity of active and silent processes in WM

Our data speak to the current debate on the nature of WM representations in the brain. Traditional models emphasize stable, persistent neural activity as the main candidate mechanism supporting WM (Fuster and Alexander, 1971; Kamiński et al., 2017). More recent, multivariate investigations point towards a more dynamic view, with the contents of WM being maintained in dynamically changing patterns of neural activity or activity-silent brain states (Rose et al., 2016; Spaak et al., 2017; Stokes, 2015; Stokes et al., 2013; Trübutschek et al., 2017; Wolff et al., 2015, 2017).

Together with our previous work (Trübutschek et al., 2017), our current results suggest that sustained neural activity and activity-silent mechanisms may accommodate different processes. Storage of information in WM need not require neural activity. Without the manipulation requirement in our task, delay-period activity vanished entirely for unseen and was only intermittent for seen targets (Trübutschek et al., 2017). Such prolonged activity-silent periods occurred less frequently in the current experiment, probably because participants tried to more actively retain information about the target location in preparation for the required mental rotation. However, even in the present setting, target-related neural activity first decayed towards chance before being reactivated by the cue.

By contrast, after the symbolic cue, once subjects were manipulating the contents of their WM, neural activity was sustained throughout the remainder of the epoch, with the representation of the response emerging while the target representation slowly faded. Importantly, we observed a similar pattern of results for unseen targets. As decodability of target location vanished, it was replaced by the emergence of the guess (i.e., pre-rotation location), that was maintained until the rise of response-related neural activity. The slightly different post-cue time courses observed for the decoding of the pre-rotation location on seen and unseen trials may not indicate any meaningful difference in the type of operation deployed by the participants, but likely reflected the differential levels of certainty with which subjects performed the mental rotation, having a clear starting point on seen trials and a more fluctuating representation on unseen trials.

Taken together, then, we propose that active and activity-silent processes make distinct contributions to WM. WM maintenance can be achieved without any accompanying neural activity via activity-silent mechanisms, but WM manipulation appears to depend on active neural firing. Recent evidence from a computational model corroborates this conclusion by demonstrating that, while short-term synaptic plasticity may support short-term maintenance, persistent neuronal activity automatically emerges from learning during active manipulation (Masse et al., 2018). Moreover, similar divisions of labor between activity-silent and activity-based brain states have recently been observed for the active selection vs. maintenance of WM contents (Quentin et al., 2018). All of these data thus lend support to the emerging view that WM is best conceptualized as an activity-induced temporary and flexible shift in the functionality of a network (i.e., dynamic coding; Stokes, 2015).

### Tracking intermediate representations during a mental rotation

A last aspect of our work that deserves attention concerns the act of mental rotation itself. Numerous behavioral and neuroimaging studies support the idea that mental rotation depends on analog spatial representations, with the initial representation progressively being rotated through intermediate positions or views. Reaction times have been found to increase in near-linear fashion with the size of the rotation angle (Cooper, 1975; Shepard and Cooper, 1986; Shepard and Metzler, 1971), and activity in spatially mapped brain areas, such as the posterior parietal cortex, has been reported to be modulated parametrically by angular distance (Gauthier et al., 2002; Jordan et al., 2001; Wager and Smith, 2003). Recordings of single-neuron activity from the motor cortex during a motor rotation task also suggest a gradual rotation of a neural population vector (Georgopoulos et al., 1989).

Our results indicate that such a transformation of neural representations is now decodable from human MEG recordings. On seen trials, following the rotation cue, average decoder estimates of target and response angle progressively moved away from the original target location towards the expected response position, seemingly passing through a series of intermediate locations. A similar transformation may also have been present for the pre-rotation location for unseen targets, though data were too noisy to support any definitive conclusions. These findings are compatible with the view that locations intermediate between the target/prerotation position and the response location were coded and represented in the brain. However, this interpretation is based on an analysis of multivariate estimates averaged across trials and participants. Isolated bursts of activity, occurring at different points in time and coding for discrete spatial positions, if averaged over many events, might also result in the apparent smooth transition we observed here (Lundqvist et al., 2016; Stokes and Spaak, 2016). Future research relying on single-trial analyses will be needed to disambiguate between these alternatives.

## Conclusion

In the wake of recent proposals of non-conscious and/or activity-silent WM, we have identified an important boundary condition: While the storage of information in WM requires neither consciousness nor persistent activity, the manipulation of WM contents is associated with both. This conclusion is at odds with the very idea of non-conscious *working* memory. We therefore propose “activity-silent short-term memory” as an alternative term for the phenomenon of long-lasting blindsight. This observation may also help reconcile current debates on the nature of WM. WM is a generic term that refers to a conglomerate of cognitive processes including attentional selection, storage, and manipulation. Active and activity-silent brain states both contribute to produce these behaviors, and an essential goal for future research will be to further disentangle their differential contribution to WM.

## Methods

### Participants

23 healthy volunteers (4 men; *M*_age_ = 23 years, *SD*_age_ = 2.5 years) with normal or corrected-to-normal vision were included in the behavioral experiment. Another 30 participants (14 men; *M*_age_ = 25.4 years, *SD*_age_ = 3.8 years) were entered in the analyses of the MEG study. In compliance with institutional guidelines, all subjects gave written informed consent prior to enrollment and received up to 80€ as compensation.

### WM task

We adapted our previous paradigm (Trübutschek et al., 2017) to probe participants’ ability to manipulate WM representations under varying levels of subjective visibility (**Figure 1**). Following a 1 s fixation period, a small, gray target square was flashed for 17 ms in 1 of 24 circular locations and subsequently masked (233 ms). Mask contrast was calibrated separately for each subject to yield ∽equal proportions of seen and unseen trials (see below). Halfway throughout a 3 s delay period, a centrally presented, symbolic cue in white ink instructed participants as to the specific task to be performed: A third of the trials, indexed by an equal sign, served as a control condition, requiring subjects to maintain and identify the position in which the target had appeared. On the remainder of the trials, participants were to mentally rotate the original target location and report this rotated position. While the uppercase letter *D* necessitated a 120° clockwise rotation (1/3 of the trials), the letter *G* indicated a 120° counterclockwise rotation (1/3 of the trials). Subjects responded by either speaking (MEG experiment; 2.5 s) or typing on a standard AZERTY keyboard (behavioral experiment; 3 s) the letter – out of a set of 24 (excluded: j, p) randomly presented in all possible locations, – corresponding to the desired position. For example, had the cue in **Figure 1** been an equal sign, participants would have had to report the letter *w*. Had it been a *D*, the correct answer would have been *b*. With the trial as shown, subjects should have indicated the letter *g*. Importantly, a location response was required even when participants had not seen the target square; in that case, they were instructed to guess the correct final position. Subjects then rated their visibility of the target on the 4-point Perceptual Awareness Scale (Ramsøy and Overgaard, 2004), using the index, middle, ring, and little finger of their right hand to operate either the number-pad keys of the computer keyboard (behavioral experiment; 2 s) or the buttons of a non-magnetic response box (Fiber Optic Response Pad, Cambridge Research Systems Ltd; MEG experiment; 2 s). To qualify as unseen (visibility = *1*), participants were to have no visual experience whatsoever of the target stimulus as well as no hunch concerning its location. All other subjective impressions were to be categorized as seen (visibility *2*, *3*, or *4*). Inter-trial intervals (ITIs) ranged between 333 and 666 ms (MEG experiment) or between 1 and 2 s (behavioral experiment). A central fixation cross was shown throughout the entire trial, and 20% target-absent catch trials were included to allow for the computation of objective measures of subjects’ perceptual sensitivity and for the isolation of brain activity specific to the target square.

### Calibration task

Participants performed a separate calibration procedure to identify the mask contrast needed for roughly equal proportions of seen and unseen targets in the WM paradigm. Trials were identical to the first part of the main experimental task (up to, and including, the presentation of the mask), but required either an immediate target localization and visibility response (behavioral experiment) or just an instantaneous visibility rating (MEG experiment). Mask contrasts were adjusted on a trial-by-trial basis with a double-staircase technique: We first divided the color spectrum between black and white into 20 equally spaced hues. Following an unseen target (visibility = *1*), mask contrast was reduced by one step on the subsequent trial, whereas it was increased by the same amount when subjects had seen the target (visibility > *1*). Initial values for the two staircases were set to RGB values of 12.75, 12.75, 12.75 and 242.5, 242.5, 242.5, respectively, and one of the two staircases was selected randomly at the beginning of each trial. In case of target-absent trials, the previous mask contrast from a randomly chosen staircase was re-used without being updated. We computed individual mask contrasts for the WM task by taking the grand average of the last four switches (i.e., from seen to unseen or vice versa) across the two staircases.

### Experimental protocol

Each experimental session began with written and verbal instructions for all tasks. Subjects then performed either 60 (behavioral experiment; 1 block) or 90 training trials (MEG experiment; 2 blocks) of the WM paradigm. In contrast to the main experiment, during this training session, the target stimulus was always visible (mask set to the lowest contrast possible) and visual feedback on localization and rotation performance was provided at the end of each trial (2.5 s): The target location, connected by a white arc to the correct response position (in green ink), was displayed. If the participant had answered incorrectly, this location was also shown in red ink. Following the training, participants completed the calibration and WM task. While the former was comprised of 125 trials (1 block) in the behavioral and 120 trials (1 block) in the MEG experiment, the latter consisted of 180 (2 blocks; 2 repetitions of each of the three rotation conditions/location) and 450 trials (10 blocks; 5 repetitions of each of the three rotation conditions/location), respectively.

### Behavioral analyses

We followed our previous approach (Trübutschek et al., 2017) to evaluate WM performance as a function of subjective visibility. Repeated-measures analysis of variance (ANOVA) was applied to three indices of objective performance: (1) Accuracy refers to that proportion of trials that falls exactly onto the correct response location and serves as a crude measure of the amount of information which can be maintained and manipulated in WM. Chance performance corresponds to 1/24 (i.e. 4.17%). (2) The rate of correct responding also reflects the quantity of information held in WM, but is more refined than accuracy alone, as it allows accounting for small errors in subjects’ ability to identify the correct response location. It was defined as the proportion of trials within ± 2 positions of the correct response location (i.e., ± 30°), leading to a chance-level of 5/24 (i.e., 20.83%). (3) As an estimate of the precision of WM representations, we computed the standard deviation of that part of the distribution of participants’ spatial responses that corresponded to genuine WM (as opposed to random guessing within the region of correct responding; Trübutschek et al., 2017). Only subjects with sufficient blindsight (i.e., *p* < .05 in a *χ*^2^-test against chance) when collapsing across all experimental conditions were included in this analysis.

### MEG acquisition, preprocessing, and decomposition

We installed participants inside an electromagnetically shielded room and recorded their brain activity continuously during the WM paradigm with a 306-channel, whole-head magnetometer by Elekta Neuromag^®^ (Helsinki, Finnland). MEG sensors were arranged in 102 triplets, comprised of one magnetometer and two orthogonal planar gradiometers, and MEG signals were acquired at a sampling rate of 1000 Hz with a hardware bandpass filter between 0.1 and 330 Hz. To allow for offline rejection of artifacts induced by eye movements and heartbeat, we monitored these bodily functions with vertical and horizontal electro-oculograms (EOGs) and electrocardiograms (ECGs). Subjects’ head position inside the MEG helmet was inferred at the beginning of each run with an isotrack Polhemus Inc. system from the location of four coils placed over frontal and mastoïdian skull areas.

We adapted Marti and colleagues’ (2015) preprocessing pipeline. First, we identified bad MEG channels visually in the raw signal and then employed MaxFilter software (ElektaNeuromag^®^, Helsinki, Finland) to (1) compensate for head movements between experimental blocks by realigning all data to the head position of the first run and (2) apply the signal space separation algorithm (Taulu et al., 2004) to suppress magnetic interference from outside the sensor helmet and interpolate bad channels. We then switched to Fieldtrip for further preprocessing (Oostenveld et al., 2011). Continuous data were first epoched with respect to target onset (i.e, −0.5 to 3.5 s). The resulting trials were downsampled to 250 Hz, and any artifacted epoch removed by means of a semi-automatic procedure: We visually inspected scatter plots of the trial-wise variance of the MEG signals across all sensors to identify and reject contaminated epochs. In a last step, we performed independent component analysis (ICA) separately for each channel type to remove any residual artifacts related to eye movements or cardiac activity: Topographies of the first 30 components were displayed for visual inspection, their time courses correlated with the EOG/ECG signals, and contaminated components subtracted from the MEG data.

Depending on the nature of the subsequent investigation, further preprocessing steps then diverged. For any univariate analysis based on evoked responses (i.e., ERFs), we only low-pass filtered the MEG signal at 30 Hz. However, to extract the spectral component of our data, we relied on unfiltered epochs: Power estimates between 1 and 99 Hz (in 2 Hz steps) were obtained by convolving overlapping segments of the data with a frequency-independent Hann taper (window size: 500 ms, step size: 20 ms). Multivariate analysis required additional downsampling of the signal to 125 Hz. After all necessary transformations and decompositions, we applied a baseline correction prior to any analysis between −200 and 0 ms.

### Estimating chance-free brain activity for unseen correct trials

To account for chance-responding on unseen correct trials, we employed a strategy developed by Lamy and colleagues (2009) and first calculated the proportion of unseen correct trials correctly responded to by chance separately for each subject:

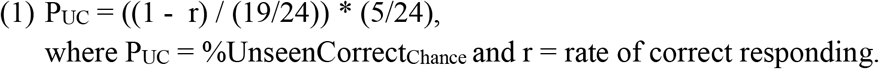

We then estimated brain activity on the unseen correct trials reflecting chance-free responding, operating under the assumption that the actual observed amplitude *A* was a linear combination of genuine blindsight and random guessing:

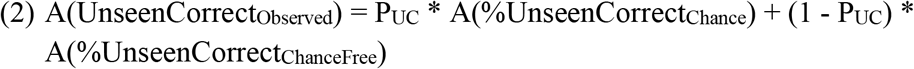

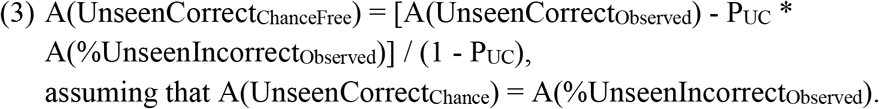

Similarly, we then reverted the process, mixing activity from seen trials with that from unseen incorrect trials, to obtain an estimate of what brain activity might have looked like under the miscategorization hypothesis.

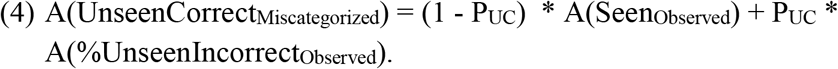

### Source reconstruction

Structural magnetic resonance (MR) scans were available for 29 of our 30 subjects, having been acquired as part of previous experiments from our lab with a 3D T1-weighted spoiled gradient recalled pulse sequence (voxel size: 1 * 1 * 1 mm; repetition time [TR]: 2,300 ms; echo time [TE]: 2.98 ms; field of view [FOV]: 256 * 240 * 176 mm; 160 slices). To identify the anatomical locations of the MEG signals in these participants, we first segmented subjects’ T1 images into gray/white matter using FreeSurfer (https://surfer.nmr.mgh.harvard.edu/) and then reconstructed the cortical, scalp, and head surfaces in Brainstorm (Tadel et al., 2011). Coregistration between the anatomical scans and the MEG data was based on participants’ head position in the MEG helmet, recorded and tracked throughout the entire experiment. Subject-specific forward models relied on analytical models with overlapping spheres. Separately for each condition and participant, we modeled neuronal current sources with a constrained weighted minimum-norm current estimate (wMNE; depth-weighting factor: 0.5). Noise covariance matrices were computed from ~5 min-long empty-room recordings, measured immediately after each individual subject. Prior to group analysis, single-trial source estimates were either (1) averaged within each subject and condition, transformed into *z*-scores relative to our pre-stimulus baseline (−0.2 – 0 s), rectified, and spatially smoothed over 5 mm, or (2), in the case of time-frequency decompositions, transformed into average power in the alpha (8 – 12 Hz) and low (13 – 20 Hz) as well as high beta (20 – 27 Hz) bands with complex Morlet wavelets (Brainstorm default parameters). We then computed the contrasts of interests and projected the resulting participant-specific source estimates on a generic brain model built from the standard template of the Montreal Neurological Institute (MNI). Group averages for spatial clusters of at least 50 vertices and thresholded at 50% of the maximum amplitude are shown for each time window under consideration (cortex smoothed at 60%).

### Multivariate pattern analysis (MVPA)

In this set of analyses, we aimed at predicting the identity and/or value of a specific categorical (i.e., visibility, accuracy) or circular (i.e., target, pre-rotation, or response location) variable (y) from single-trial brain activity *(X)* separately for each participant and time point. Relying on the Scikit-Learn package (Pedregosa et al., 2011) for MNE 0.15 (Gramfort, 2013; Gramfort et al., 2014), we therefore adapted the pipeline developed by King and colleagues (2016) to (1) fit a linear estimator *w* to a training subset of *X*(*X*_train_) to isolate the topographical patterns best differentiating our experimental conditions, (2) predict an estimate of *y* (*ŷ*) from a test set (*X*_test_), and (3) compare the resulting predictions to the true value of *y* either for the entire set of labels *(score(y, ŷ))* or a specific subset *(subscore(y, ŷ)*).

Here, two main classes of estimators were used: A linear support vector machine (SVM) was employed in the case of categorical, and a combination of two ridge regressions in the case of circular data. Whereas the former was set to generate a continuous output in the form of the distance between the hyperplane (*w*) and the respective sample of *y*, the latter first separately fit the sine *(sin(y))* and cosine *(cos(y))* of the spatial position in question and then estimated an angle from the arctangent of the individual predictions (ŷ = *arctan2(ŷ_sin_,ŷ_cos_*)). To increase the number of instances available for each circular label, we averaged neighboring spatial locations (effectively reducing the number of positions from 24 to 12). Prior to model fitting, all channel-time features *(X)* were z-score normalized, and, for any analysis involving SVMs, a weighting procedure applied to counteract the effects of potential class imbalances. All other model parameters were left with their Scikit-Learn default values.

To avoid overfitting, we embedded this sequence of analysis steps in a 5-fold, stratified cross-validation procedure: For non-independent training and test sets, estimators were iteratively fitted on 4/5^th^ of the data (*X*_train_) and generated predictions for the remaining 1/5^th^ (*X*_test_). By contrast, when generalizing from one task to the other (i.e., no-rotation to rotation condition), estimators from each training set were directly applied to the entire test set and the respective predictions averaged. Within the same cross-validation loop, we also evaluated time generalization (King and Dehaene, 2014): Each estimator was first trained at time *t* and then tested at all other time points, resulting in a square matrix of training time x testing time. As such, this temporal generalization analysis permits an interrogation of the durability and stability of patterns of brain activity.

We summarized within-participant, across-trial decoding performance of categorical data with the area under the curve (AUC), presenting an unbiased measure of the true-positive rate as a function of the false-positive rate (range: 0 – 1; chance = 0.5). Two different summary statistics were used for circular decoding: (1) For non-directional analyses, the mean absolute difference between the predicted (*ŷ*) and actual angle (*y*) across all trials was first computed (range: 0 – π; 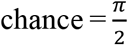), and this “error metric” was then transformed into an “accuracy score” 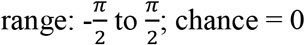. (2) In contrast, the probability distribution of the signed difference betweenŷ and an actual location was retained for directional analysis (i.e., tracking the rotation itself). The resulting, continuous angular distance estimates were then assigned to 1 of 24 evenly spaced bins (discontinuous; range: [-π, : π/24 : π]) and the probability of a given estimate falling within the range of a given bin was calculated across trials.

### Statistical analysis

All statistics reported in the text refer to group-level analyses. In the case of ERF and frequency data, we (1) performed cluster-based, non-parametric *t*-tests with 1,000 Monte Carlo permutations to identify significant spatio-temporal differences between experimental conditions, while simultaneously correcting for multiple comparisons (Maris and Oostenveld, 2007), and (2) additionally present uncorrected outcomes of non-parametric signed-rank tests for follow-up analyses of amplitude/power differences in time courses (*p*_uncorrected_ < .05). We again relied on the above cluster-based permutation analysis to assess multivariate decoding performance (i.e., categorical data: AUC > 0.5; circular data: rad > 0; 5000 permutations). Temporal averages over five a-priori time bins, corresponding to an early perceptual period (0.1 – 0.3 s), the P3b time window (0.3 – 0.6 s), the maintenance period before (0.6 – 1.76 s) and after the cue (1.76 – 3.26 s), as well as the response (3.26 – 3.5 s), are also provided. Bonferonni correction was applied to these a-priori analyses to correct for multiple comparisons (*p*_corr_ < .05/5). When appropriate, we present circular statistics and computed Bayesian statistics based on two-or one-sided *t*-tests (*r* = .707; Rouder et al., 2009).

## Acknowledgements

This work was funded by INSERM, CEA, Collège de France, ERC, and Fondation Roger de Spoelberch. D.T. was funded by a graduate fellowship from the Ecole des Neurosciences de Paris (ENP) and Fondation Schneider Electric. We gratefully acknowledge Valentina Borghesani, Pedro Pinheiro Chagas, and Fosca Al Roumi for their invaluable daily support and stimulating discussion and specifically thank Theofanis I. Panagiotaropoulos for helpful comments on a previous version of this manuscript.

## Author contributions

Conceptualization, D.T., S.M., and S.D.; Methodology, D.T., S.M., H.U., S.D.; Formal analysis, D.T.; Investigation, D.T.; Resources, S.D.; Writing – Original Draft, D.T.; Writing – Review and Editing, D.T., S.M., H.U., and S.D.; Visualization – D.T.; Project Administration - D.T.; Funding Acquisition – S.D.

## Supplementary Figures

**Supplementary Figure 1.**
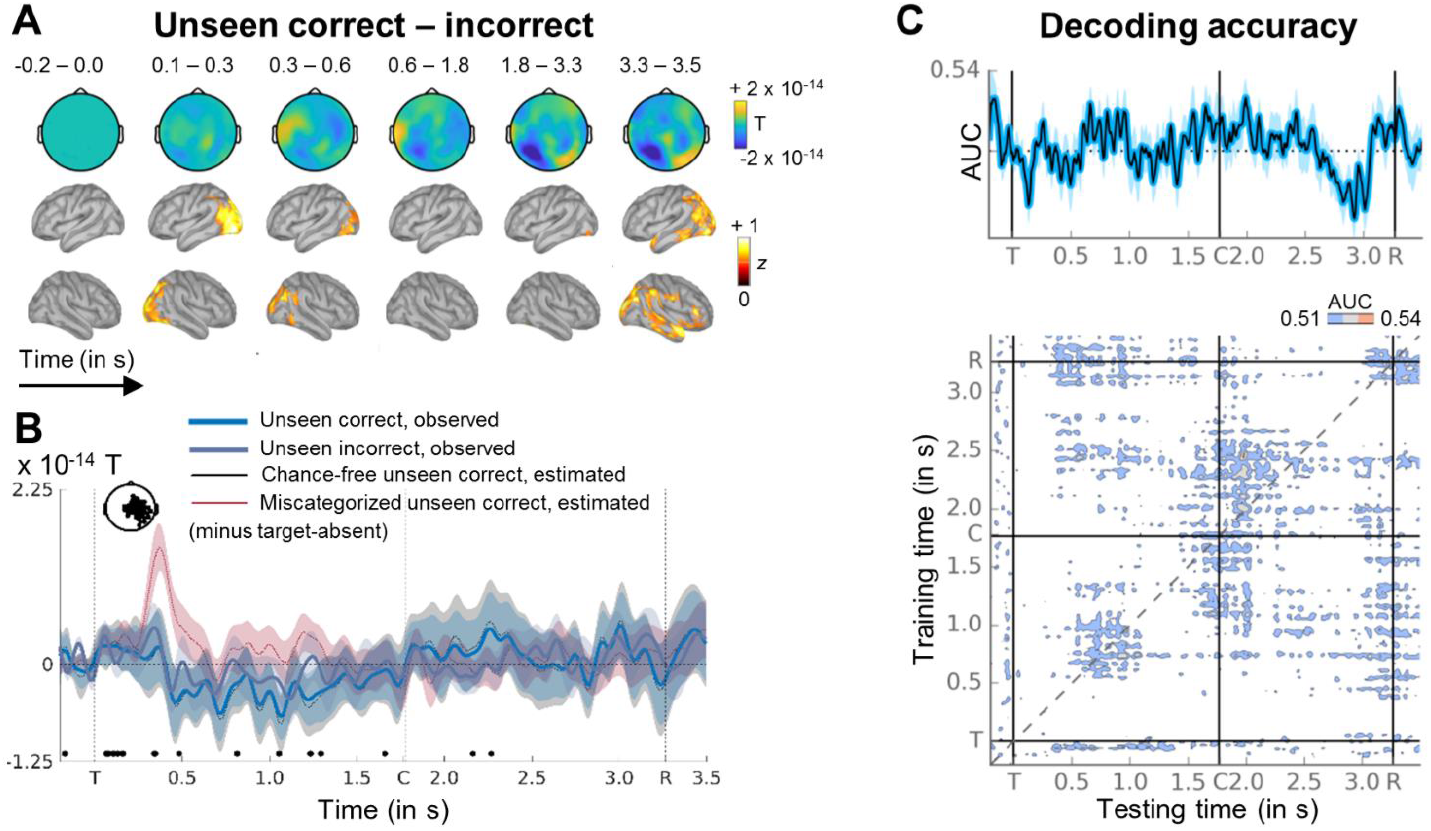
No signatures of conscious processing on the unseen correct trials. **(A)** Sequence of brain activations (−0.2 – 3.5 s) evoked by non-consciously perceiving the target in both tasks in sensor (top) and source space (bottom). Each topography depicts the difference in amplitude between unseen correct and unseen incorrect trials averaged over the time window shown (magnetometers only). Sources reflect z-scores of absolute difference with respect to a pre-stimulus baseline. **(B)** Average time courses (−0.2 – 3.5 s) of unseen correct (light blue) and unseen incorrect (dark blue) trials in that subset of magnetometers having shown a significant difference in amplitude between seen and unseen targets. Black trace reflects brain activity on the unseen correct trials after having been corrected for chance-responding. Red time course illustrates what the signal on the unseen correct epochs should have looked like, had the miscategorization hypothesis been true. Shaded area denotes standard error of the mean (SEM) across subjects. Significant differences between unseen correct and incorrect epochs are depicted with the thick, black line (two-tailed Wilcoxon signed-rank test, uncorrected). Vertical dotted lines index onset of the target (T), symbolic cue (C), and response (R) screens. For display purposes only, data were lowpass-filtered at 8 Hz. (C) (Top) Average time course of diagonal decoding of accuracy on the unseen trials (i.e., unseen correct vs. unseen incorrect). Horizontal, dotted line represents chance level at 50%. (Bottom) Temporal generalization matrix of the same accuracy decoder. Each horizontal row in the matrix corresponds to an estimator trained at time *t* and tested on all other time points t’. The diagonal gray line demarks classifiers trained and tested on the same time points (i.e., the diagonal estimator shown on top). In both plots, vertical lines mark onset of the target (T), symbolic cue (C), and response (R) screens. Only for display purposes, data were smoothed with a moving average of 5 samples (i.e., 40 ms). AUC = area under the curve.

**Supplementary Figure 2.**
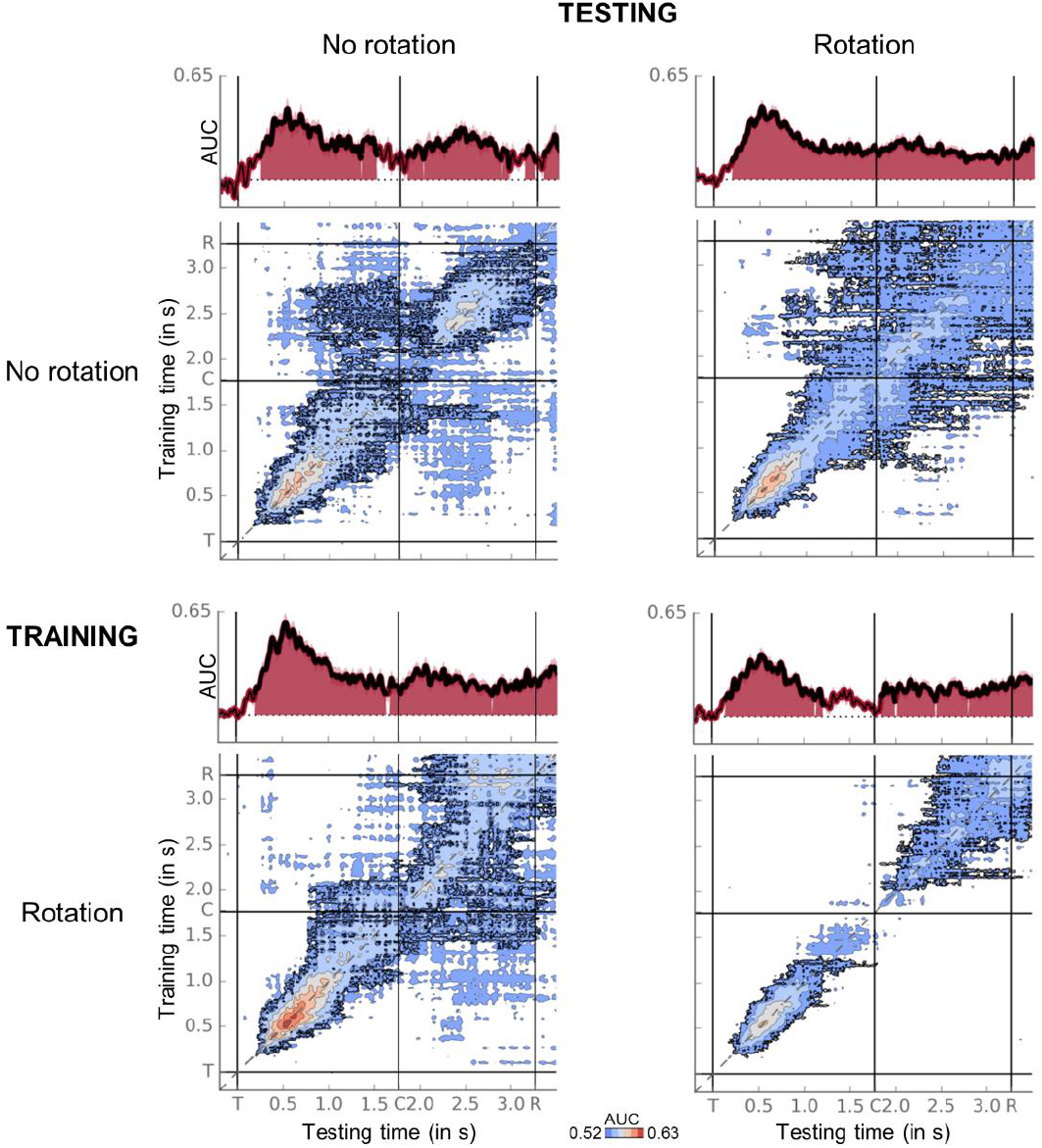
Conscious perception entails similar neural dynamics in both tasks. Temporal generalization matrices (bottom) for decoding of visibility category (i.e., seen vs. unseen) as a function of training and testing task (i.e., no rotation vs. rotation). In each panel, a classifier was trained at every time sample (y-axis) and tested on all other time points (x-axis). The diagonal gray line demarks classifiers trained and tested on the same time sample. Event markers (i.e., vertical/horizontal lines) denote onset of the target (T), cue (C), and response (R) screens. Time courses of diagonal decoding are shown on top. Black outlines in matrix plots and thick lines/shaded areas in time courses show periods of significant decoding (cluster-based permutation test, twotailed except for diagonal). For display purposes, data were smoothed using a moving average with a window of 5 samples (i.e., 40 ms). AUC = area under the curve.

**Supplementary Figure 3.**
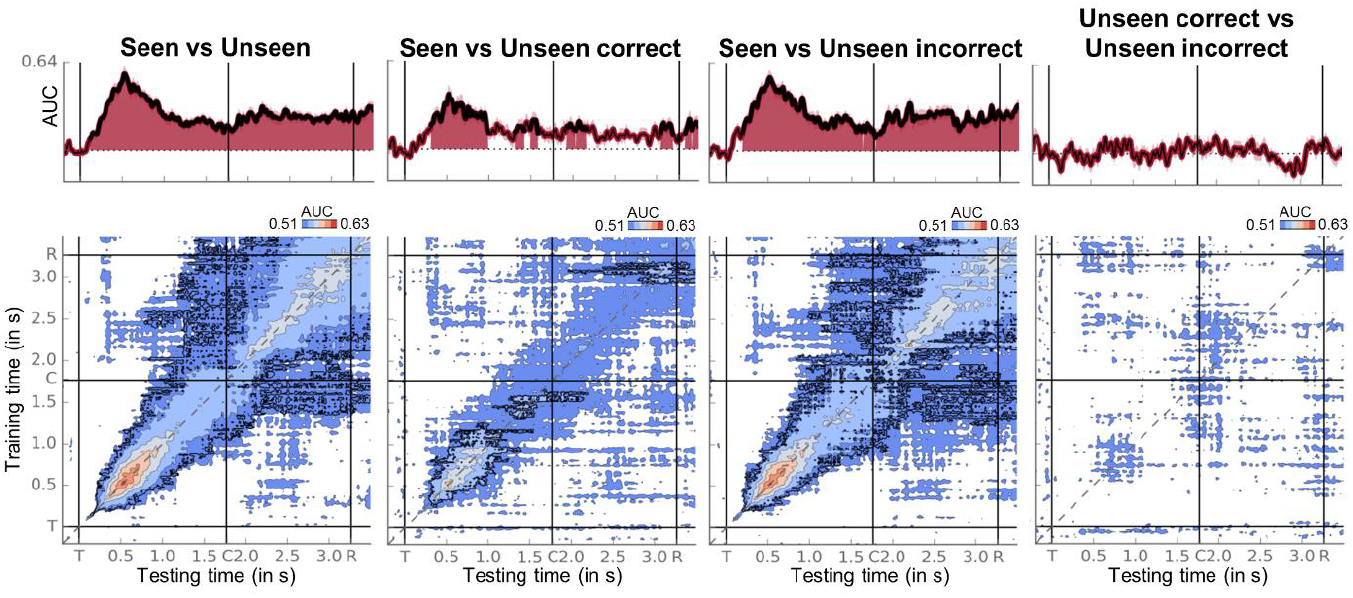
Comparing visibility to accuracy decoder. Each panel displays the generalization matrix (bottom) and time course of diagonal decoding (top) of a specific visibility or accuracy estimator. Horizontal, dotted line in time course represents chance level at 50%. Each horizontal row in the matrix corresponds to an estimator trained at time *t* and tested on all other time points *t*’. The diagonal gray line demarks classifiers trained and tested on the same time points (i.e., the diagonal estimator shown on top). In both plots, vertical lines mark onset of the target (T), symbolic cue (C), and response (R) screens. Thick lines/shaded areas as well as black outlines denote above-chance decoding as assessed by a cluster-based permutation test (two-tailed, with the exception of the diagonal). Only for display purposes, data were smoothed with a moving average of 5 samples (i.e., 40 ms). AUC = area under the curve.

**Supplementary Figure 4.**
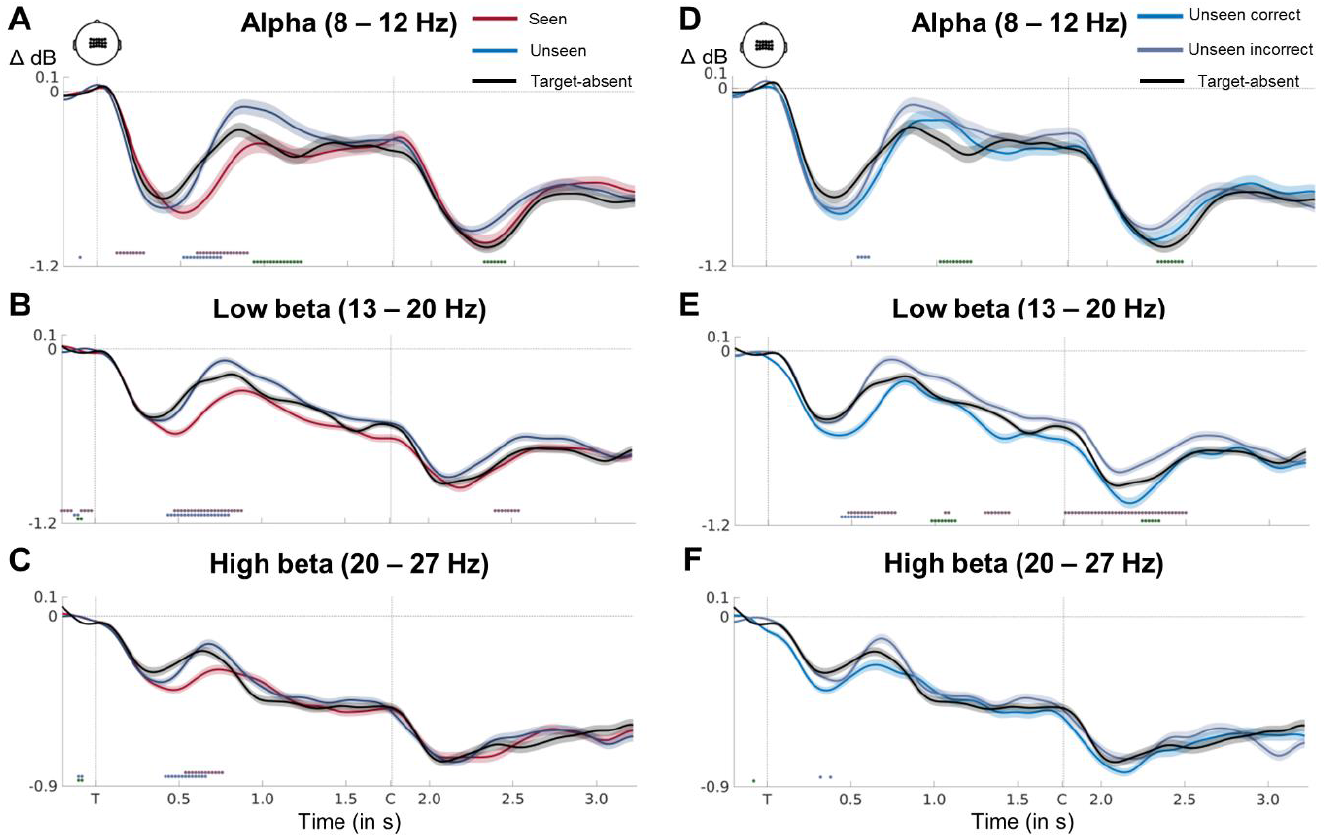
Average time courses of alpha, low beta, and high beta power. Time courses of average alpha (8 – 12 Hz; **A**), low beta (13 – 20 Hz; **B**), and high beta (20 – 27 Hz; C) band activity in a group of central sensors as a function of visibility and target presence. Shaded area demarks standard error of the mean (SEM) across subjects. Thick lines represents significant difference in power between conditions (red = seen vs. unseen; blue = seen vs. target-absent; green = unseen vs. target-absent; two-tailed Wilcoxon signed-rank test across subjects, uncorrected). Vertical line demarks onset of target (T) and cue (C) screens. (**D-F**) Same as in (**A-C**), except for unseen correct and unseen incorrect trials. Color code for significant differences is as follows: red = unseen correct vs. unseen incorrect, blue = unseen correct vs. target-absent, green = unseen incorrect vs. target-absent.

**Supplementary Figure 5.**
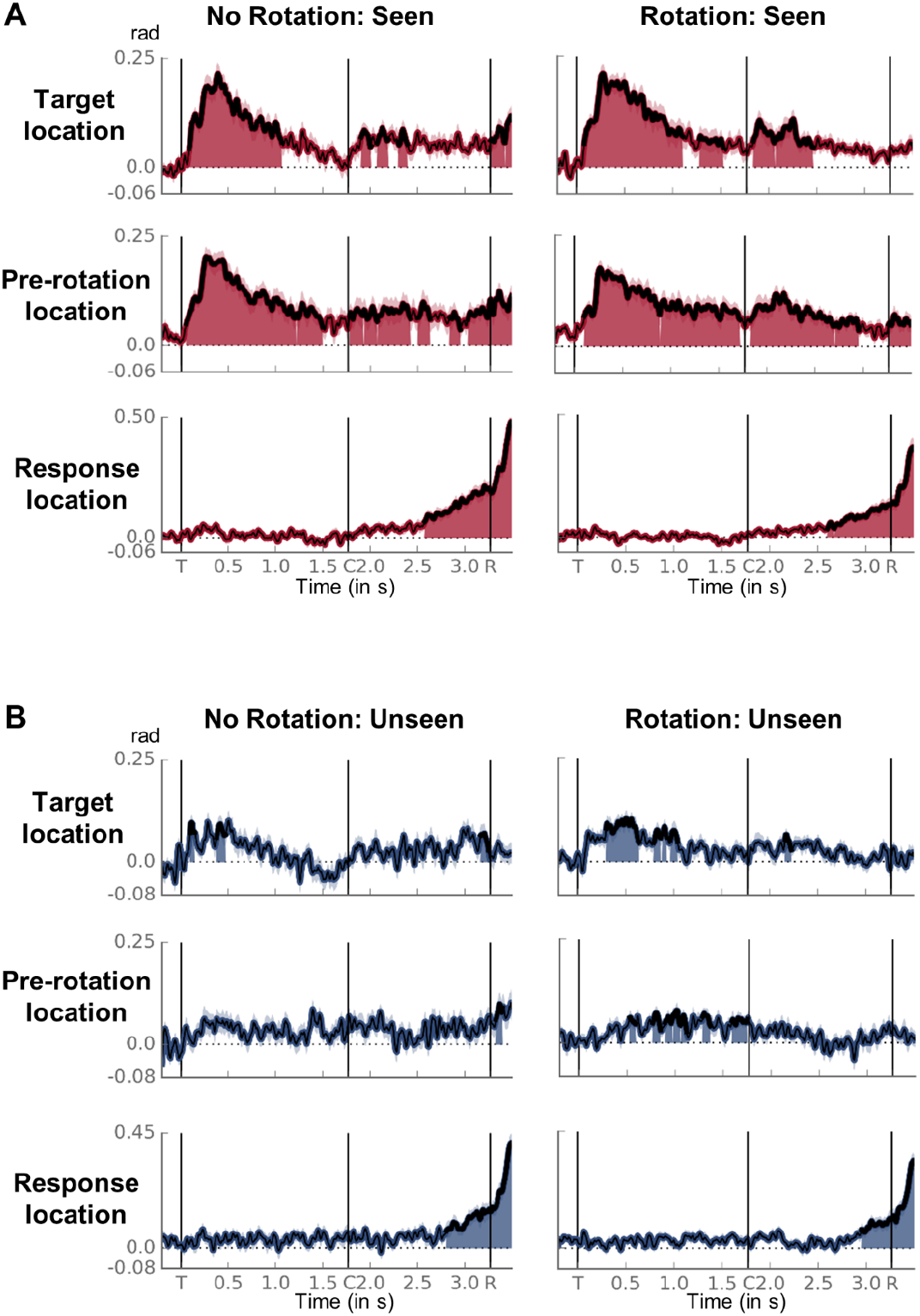
Tracking a mental rotation on seen and unseen trials in the rotation and no-rotation task. **(A)** Time courses of average decoding of target location (top), pre-rotation location (middle) and response location (bottom) on seen trials as a function of task (i.e., no rotation vs. rotation). Thick lines and shaded areas represent above-chance performance as assessed by a one-tailed cluster-based permutation test. Horizontal dotted lines index chance. Event markers denote the onset of the target (T), cue (C), and response (R) screens. For illustration purposes, data were smoothed with a moving average of 5 samples (i.e., 40 ms). **(B)** Same as in (A), but for unseen trials.

**Supplementary Figure 6.**
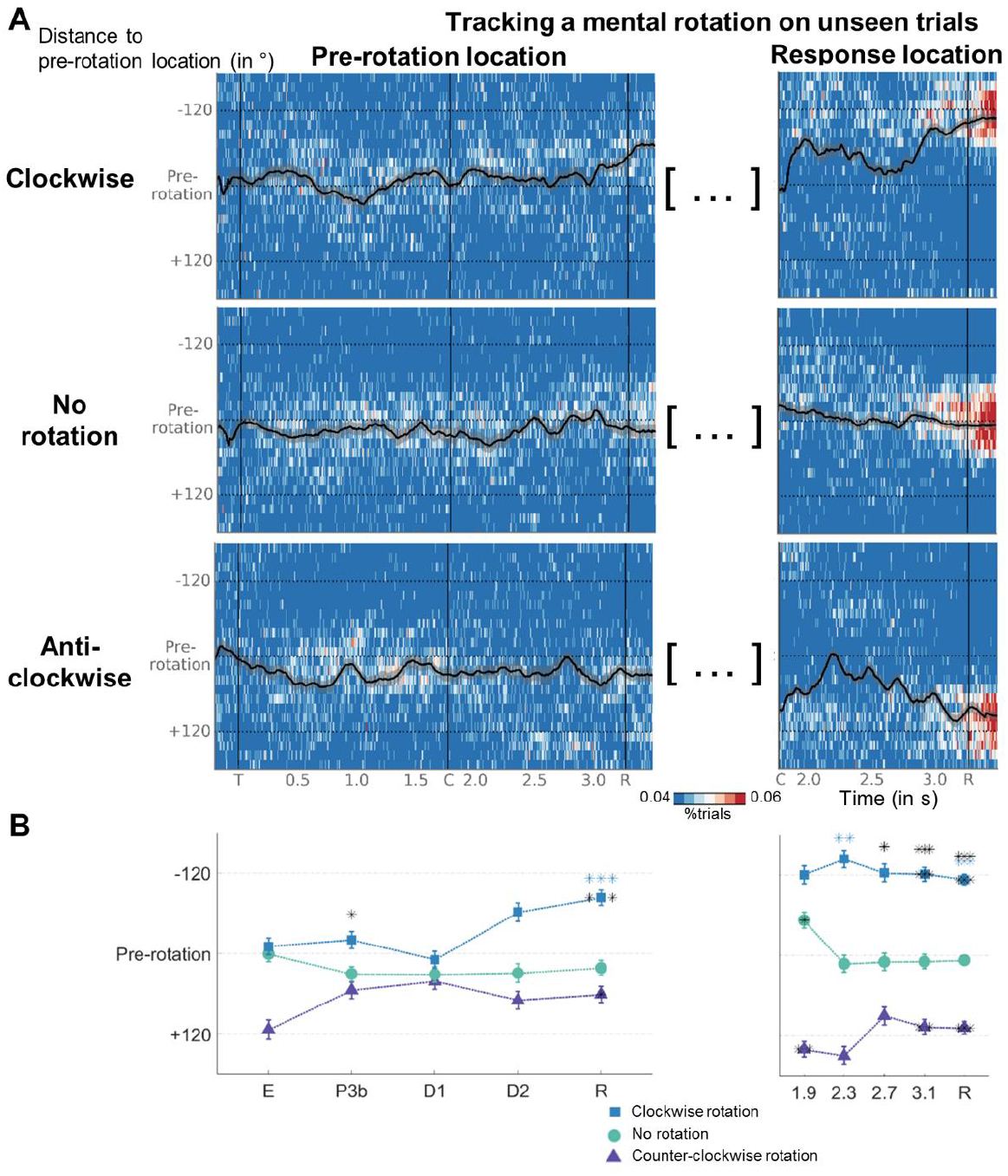
Tracking a mental rotation on unseen trials. **(A)** (Left) Time courses of probability density distributions of the angular distance between the estimates of a decoder trained with pre-rotation angle and actual pre-rotation location are shown as a function of rotation condition. For display purposes only, data were smoothed with a moving average of 12 samples (i.e., 96 ms). Overlaid black line illustrates the evolution of the circular mean of the individual distributions (also smoothed). Shaded area reflects circular standard variation across subjects. Vertical event markers denote the onset of the target (T), cue (C), and response (R) screens, horizontal markers index correct response positions after rotation. (Right) Same as in the left panels, except for angular distance between the estimates of a decoder trained with response angle and actual pre-rotation location. **(B)** Circular means of the above distributions as a function of rotation condition and time bin (i.e., E = 100 – 300 ms, P3b = 300 – 600 ms, D1 = 0.6 – 1.76 s, D2 = 1.76 – 3.26 s, R = 3.26 – 3.5 s). Error bars reflect circular standard deviation. Asterisks inside markers denote significant deviation from mean direction of 0 (as assessed by a circular equivalent of a one-sample t-test), asterisks on top significant differences in median direction between conditions (as assessed by a circular equivalent to the Kruskal-Wallis test; black = clockwise vs. counter-clockwise; blue = clockwise vs. no rotation; violet = counter-clockwise vs. no rotation). * *p* < .05, ** *p* < .01, *** *p* < .001.

